# Dynamic, behavior-dependent interactions between dorsal striatal dopamine and glutamate release predict cognitive flexibility and punishment resistant cocaine use

**DOI:** 10.1101/2025.08.11.669720

**Authors:** David M Bortz, Karim Obaidi, Aubrey Deldin, Doug J Weber, Mary M Torregrossa

## Abstract

Cognitive inflexibility covaries with substance use disorder (SUD) risk. To determine if there is a neural relationship between these phenomena, glutamate and dopamine release in the dorsomedial (DMS) and dorsolateral (DLS) striatum were measured as rats performed a discrimination and strategy switching test. Elevations in glutamate release, with reductions in dopamine, at trial initiation (DLS) and prior to choice (DMS and DLS) predicted fast strategy switching and punishment sensitive cocaine seeking. Elevations in DLS and DMS dopamine release at these respective timestamps predicted slow switching and punishment resistance. Orbitofrontal cortex and intralaminar thalamus were significant contributors to DLS and DMS glutamate release, but their relative contributions differed between rats that were fast or slow strategy switchers, and in how they affected behavior. As such, these data describe a neural signature of flexibility and associated circuitry that could be used to predict and treat SUDs in humans.

## Introduction

Drug-taking despite threat of punishment or consequences is a hallmark of substance use disorders (SUDs) and may arise from the development of inflexible drug-seeking behaviors that can be triggered by environmental cues and contexts^1^. The presence of general cognitive inflexibility is also a trait marker that can predict the development and severity of substance use in rodents and non-human primates^2–5^. Family history of substance use is associated with deficits in cognitive flexibility performance, among other cognitive measures^2^. Thus, we hypothesized that a common neural mechanism could contribute to both cognitive inflexibility and risk for punishment resistant substance use.

We and others have shown that glutamate (Glut) and dopamine (DA) release in the dorsomedial striatum (DMS) is involved in the regulation of goal-directed, flexible behavior, and that the dorsolateral striatum (DLS) is involved in the initiation and execution of well-learned, stable behavior. DA release in DMS is necessary for strategy switching specifically^6^ and cognitive flexibility, in general^7,8^, and mice self-stimulating dopamine terminals in DMS show improved reversal learning^9^. Glut release in DMS from basolateral amygdala has also been shown to bidirectionally regulate goal-directed, flexible behavior^10^, with DMS activity being necessary for flexible adaptation of behavior to changing reward contingencies^11^. In contrast, DA^12^ and Gut^13,14^ release in the DLS increase prior to the execution of a well-learned action, coinciding with a necessary increase in DLS activity^15^. Furthermore, DA release in the DLS develops and strengthens over time in response to drug-related cues^16^. Our lab demonstrated that the development of outcome-insensitive, punishment resistant cocaine use coincides with a reduction in calcium-indicated neural activity in the DMS, likely due to reduced glutamate input, and increased DLS DA release to cocaine-paired cues across training^17^.

Despite the known interaction between substance use phenotypes, cognitive inflexibility, and Glut/DA dynamics in the DMS and DLS, no studies have measured how DMS and DLS DA and Glut interact during tests of cognitive flexibility or determined if those measures predict punishment resistant drug taking within the same animal. Thus, we took advantage of natural variation in rat cognitive flexibility, as assessed by a strategy switching test, to determine if poor cognitive flexibility was associated with the propensity to develop punishment resistant cocaine use across two different self-administration schedules. Furthermore, we used multi-channel fiber photometry to simultaneously measure Glut and DA release in the DMS and DLS of the same rat as they performed a discrimination paradigm and then a strategy switching test. We determined if neural signatures of cognitive flexibility could be identified and if those neural signatures could predict punishment resistant cocaine use. We found that poor performance on the strategy switching test predicted resistance to punishment, independent of self-administration schedule. We also found that coordinated fluctuations in DMS and DLS Glut and DA at key points during discrimination and strategy switching predicted both strategy switching performance and sensitivity or resistance to punishment. Finally, in a separate set of rats, we used pathway-specific designer receptors exclusively activated by designer drugs (DREADDs) to inhibit known Glut inputs to the dorsal striatum to determine the source of the measured Glut signal and whether it was necessary for strategy switching or punished cocaine taking.

## Results

### Photometry recordings

Male(N=10) and female(N=12) rats were infused with viral vectors that contain photo-emitting biosensors to detect DA and Glut into the DMS of one hemisphere and the DLS of the other hemisphere, within the same rat. All rats were verified to have sufficient presence of GRABDA(yellow) and igluSnFR(green) around the photometry fibers in the DMS and DLS (Fig. 1a).

**Figure 1.**
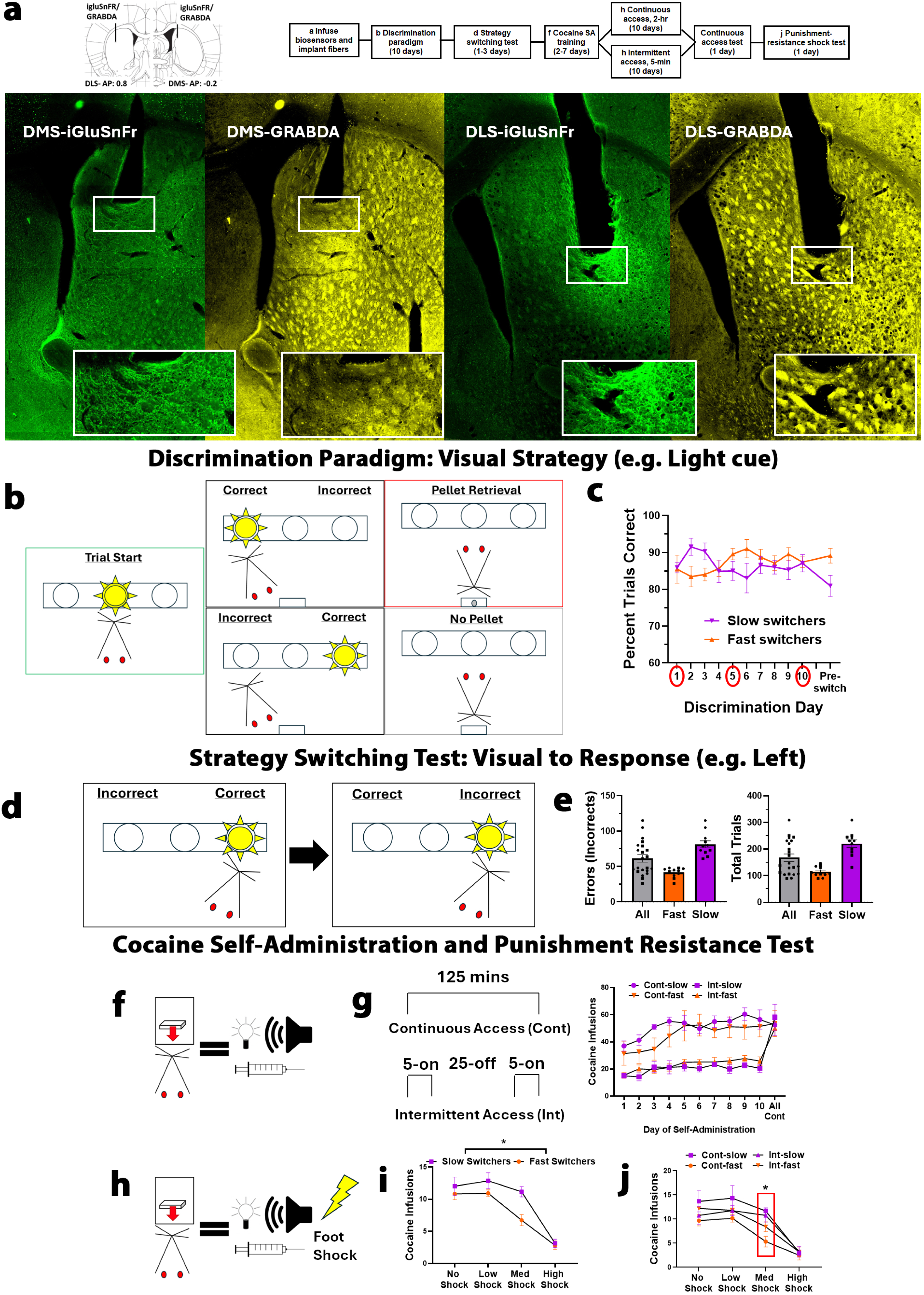
Experiment timeline, histology, and behavioral results: **a)** Representative images with closeup insets (white rectangle) of the recording zones in the DMS (left) and DLS (right) show yellow (DA) and green (Glut) immunofluorescence around the fiber track. **b)** All rats initially learned a visual strategy. **c)** Photometry recordings were performed on days 1, 5, and 10 (red circles). Fast switchers displayed a typical learning curve such that their performance peaked on day 6 (91% correct) and remained steady throughout the remaining days. Slow switchers peaked on day 2 (92%) but then declined after that and remained steady for the last few days. Slow switchers got more trials correct on days 2 (P=0.04) and 3 (P=0.04). Fast switchers got more trials correct during the 20 “pre switch” trials (89%) compared to slow switchers (81%; P=0.03). **d)** On day 11, the rewarded strategy switched from visual to a response strategy. **e**) Male (squares) and female (circles) rats were split into two groups using a median split of errors. Fast switchers (orange bar) took fewer trials (114.6±5.9) and committed fewer errors (41.6±2.1), compared to slow switchers (purple bar, trials: 220.4±14.0, errors: 81.4±5.0; both P’s < 0.0001). **f)** Rats pressed a lever (FR1) for an infusion of cocaine (0.5 mg/kg) paired with a light/tone cue. **g)** Rats were given access to cocaine continuously for a 125-minute session (continuous access), or intermittently (5-minutes on, 25 minutes off) for a 125-minute session (intermittent access) across 10 days. **h)** On day 12, cocaine infusions were paired with a foot shock. **i)** Slow switchers took more punished cocaine infusions overall (no shock: 12.0±1.4, low shock: 12.9±1.2, medium shock: 11.1±0.8, High shock: 3.1±0.6) compared to fast switchers (no shock: 10.8±0.9, low shock: 10.9±0.5, medium shock: 6.7±0.9, High shock: 2.8±0.7), **j)** particularly at the medium shock intensity where intermittent access-slow (10.8±1.4, P=0.046) and continuous access slow (11.7±0.7, P=0.0006) both took significantly more punished cocaine infusions than the continuous access-fast rats (5.3±1.1).

### Strategy switching performance was normally distributed

Following surgery, rats learned to nose-poke in an operant chamber using a visual cue to receive a sugar pellet reward (pre-training, Fig.S1). Next, rats performed a complex discrimination paradigm across 10 days (Fig. 1b). Rats discriminated between 2 active ports using a visual cue (visual strategy), while ignoring the position of the cue (response strategy; Fig.1c). On the 11^th^ day, the rats performed a strategy switching test requiring them to inhibit responding via the previously rewarded strategy (visual) and learn the newly rewarded strategy (response). We quantified the number of errors and total trials that it took the rats to reach the performance criterion of 10 consecutive trials correct using this newly rewarded response strategy. Performance was normally distributed and featured a range of responses (Fig. 1e). Males (N=10, 58.2±6.3) and females (N=12, 64.2±7.9) were part of the same interval (95%CI males:44.0-72.4, females:46.8-81.6), so sexes were combined for these studies. We then performed a median split based on errors such that rats that had greater than 61.5 errors (N=6 males, 5 females) were qualified as “slow switchers” and rats with fewer than 61.5 errors (N=4 males, 7 females) were qualified as “fast switchers” (Fig. 1e). The number of pre-training days or the presence or absence of a side bias did not predict strategy switching performance (Fig.S1c-d).

### Slow strategy switching predicts cocaine punishment resistance

After the strategy switching test had concluded, all rats were implanted with jugular vein catheters and trained to self-administer cocaine. For each lever press (FR1) rats received a light/tone cue and a 0.5 mg/kg intravenous infusion of cocaine (Fig. 1f). Slow switchers reached self-administration criterion (see methods) in fewer days than fast switchers (Fig.S1e). Rats were then given continuous or intermittent access to cocaine for 10 days. Rats with continuous access to cocaine took more cocaine infusions per day (P<0.0001) due to the nature of the continuous access schedule, but intermittent access rats took more cocaine per minute (P<0.0001; Fig.S1f). On the 11^th^ day all rats were given continuous access to cocaine and they took comparable numbers of cocaine infusions (mean range 50 to 58 infusions across all groups, Fig. 1g). On day 12, all rats performed a punishment resistance test (Fig. 1h), where cocaine infusions were paired with a foot shock that increased in intensity every 10 minutes on a tenth-log_10_ scale from 0.13 to 0.79 milliamps^18–20^. Slow switchers took significantly more punished cocaine infusions overall compared to fast switchers (F(1,48)=11.29, P=0.0015, Fig. 1i), but this effect was most apparent during the medium shock intensities (P=0.0011; Fig. 1j). This effect was independent of self-administration schedule in the slow switchers (F(3,56)=6.44, P=0.0008). Intermittent access in the fast switcher group did produce an intermediate phenotype, elevating punished cocaine infusions earned relative to the continuous-fast group (P=0.06), and was no longer significantly reduced relative to the two slow switcher groups (p’s>0.2), similar to what’s been shown previously^18,21,22^.

### Discrimination paradigm: Fast switchers exhibited elevated DLS Glut signaling during the discrimination paradigm

To determine if DMS and DLS Glut and DA release measures would be different during performance of the discrimination paradigm in rats that went on to be fast and slow switchers, we performed photometry recordings on discrimination days 1, 5, and 10. We aligned the signal to five separate task-relevant behavioral timestamps: when the rat initiates a trial (Trial Start), a correct or incorrect choice, and pellet retrieval or not (No Pellet; see Fig. 1b). Rats that went on to be fast switchers had significantly different Glut and DA dynamics in the DLS at trial start on day 1 (F(3,60)=3.6, P=0.018, Fig. 2a) and day 5 (F(1,19)=4.8, P=0.041, Fig. 2c), compared to slow switchers. By quantifying area under the curve for each release measure, we determined that fast switchers had elevated DLS Glut on Day 1 (P=0.015, Fig. 2b) and Day 5 (P=0.006, Fig. 2d), relative to slow switchers. Another interesting element of the data was not only measuring them individually, but in relation to each other. Thus, we also quantified Glut release in relation to DA release within the DLS and DMS. Interestingly, this relative measure was often more predictive of future behavior than either measure individually (see below). Fast switchers developed a greater difference between Glut and DA in DLS that was present on Day 1 (P=0.12) and reached significance by Day 5 (P=0.016). Note that these differences in signaling were observed between groups despite no significant differences in behavioral performance (Fig. 1c). By Day 10, the signaling differences were no longer apparent, as the slow switchers also displayed elevated Glut and high Glut-DA ratios in the DLS at trial start (Fig. 2e-f).

**Figure 2.**
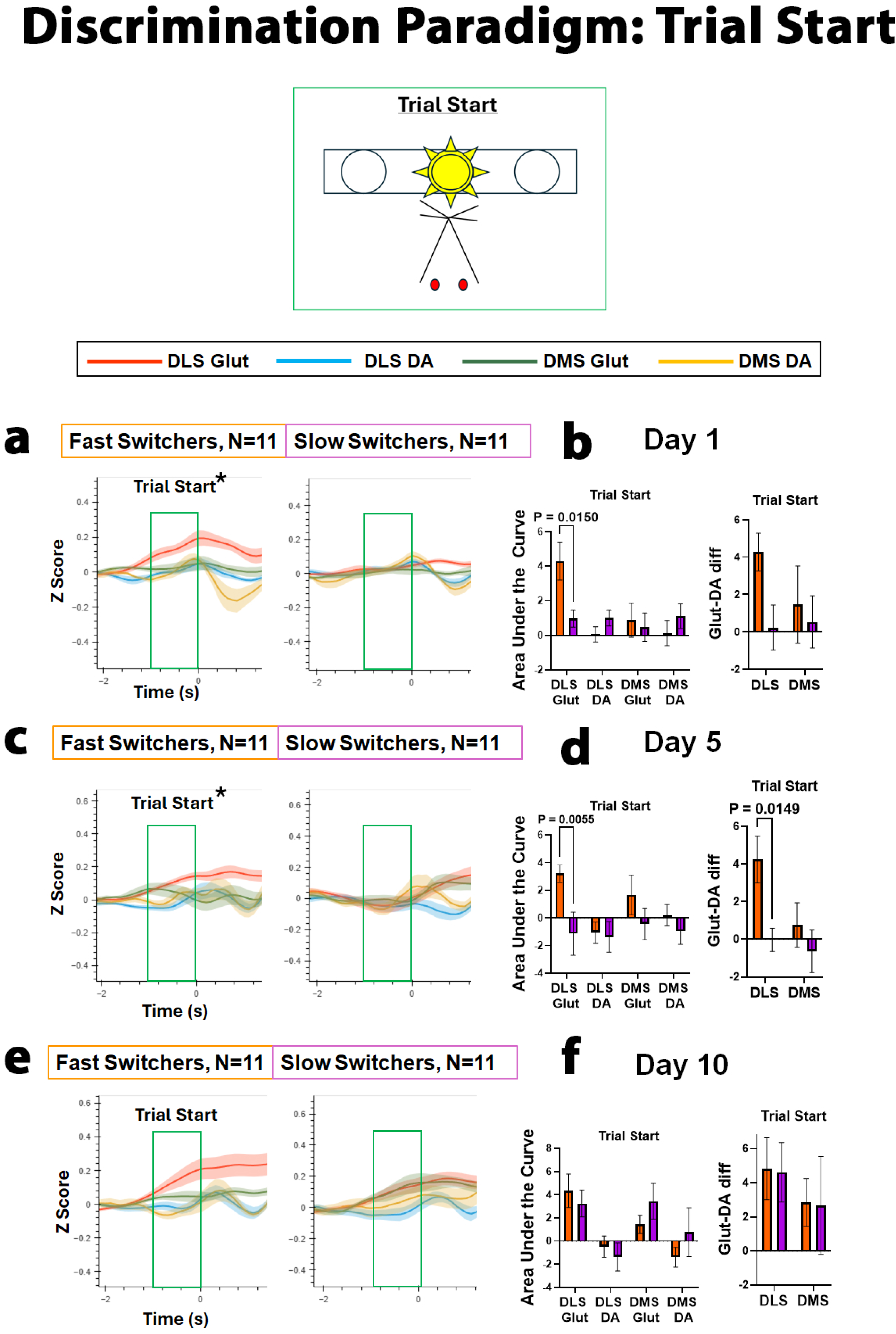
Fast switchers show elevated DLS Glut at trial start: This figure depicts mean tracings aligned to trial initiation (Trial Start) of DLS Glut (red), DLS DA (blue), DMS Glut (green), and DMS DA (yellow) release across fast (left, N=11) and slow (right, N=11) switchers. Area under the curve (Mean±SEM) for 1 second prior to trial start (as indicated by the green box) for each trace was quantified and compared across fast (orange) and slow (purple) switchers, as well as the difference between Glut and DA release in the DLS and DMS. **a,b**) On day 1 of the discrimination paradigm, rats that went on to be fast switchers showed elevated DLS Glut (4.3±1.1) as they initiated the trial, compared to slow switchers (1.0±0.5). There were no differences in the other three measures. **c,d**) This effect was also present on day 5 (DLS Glut-fast: 3.2±0.6, slow: - 1.1±1.6). While there were no differences between groups in DLS DA (fast:-1.1±0.8, slow:-1.4±1.1), fast switchers demonstrated a significant elevation in DLS Glut relative to DLS DA (fast: 4.2±1.2, slow:-0.04±0.6). There were no differences in DMS Glut or DA. **e,f)** By day 10 all measures normalized across rats.

When we aligned the signal to when the rat got a choice correct and then received a sugar pellet, we also detected differences between rats that went on to be fast and slow switchers. On Day 1, the rats that went on to be fast switchers had significantly different Glut-DA dynamics in DMS and DLS at pellet retrieval (F(1,20)=9.3, P=0.006, Fig. 3a). Area under the curve analysis found that this was driven by an elevation in DLS Glut in the fast switchers (P=0.026, Fig. 3b). On day 5, fast switchers were significantly different from slow switchers prior to a correct choice (F(1,19)=4.9, P=0.039, Fig. 3c), which was driven by elevations in DLS Glut in the fast switchers (P=0.043, Fig. 3d). Again, by day 10, fast and slow switchers showed similar patterns of dorsal striatal signaling (Fig. 3e-f). There were no differences prior to or following an incorrect choice (no pellet) across any of the 3 discrimination days (Fig. S2).

**Figure 3.**
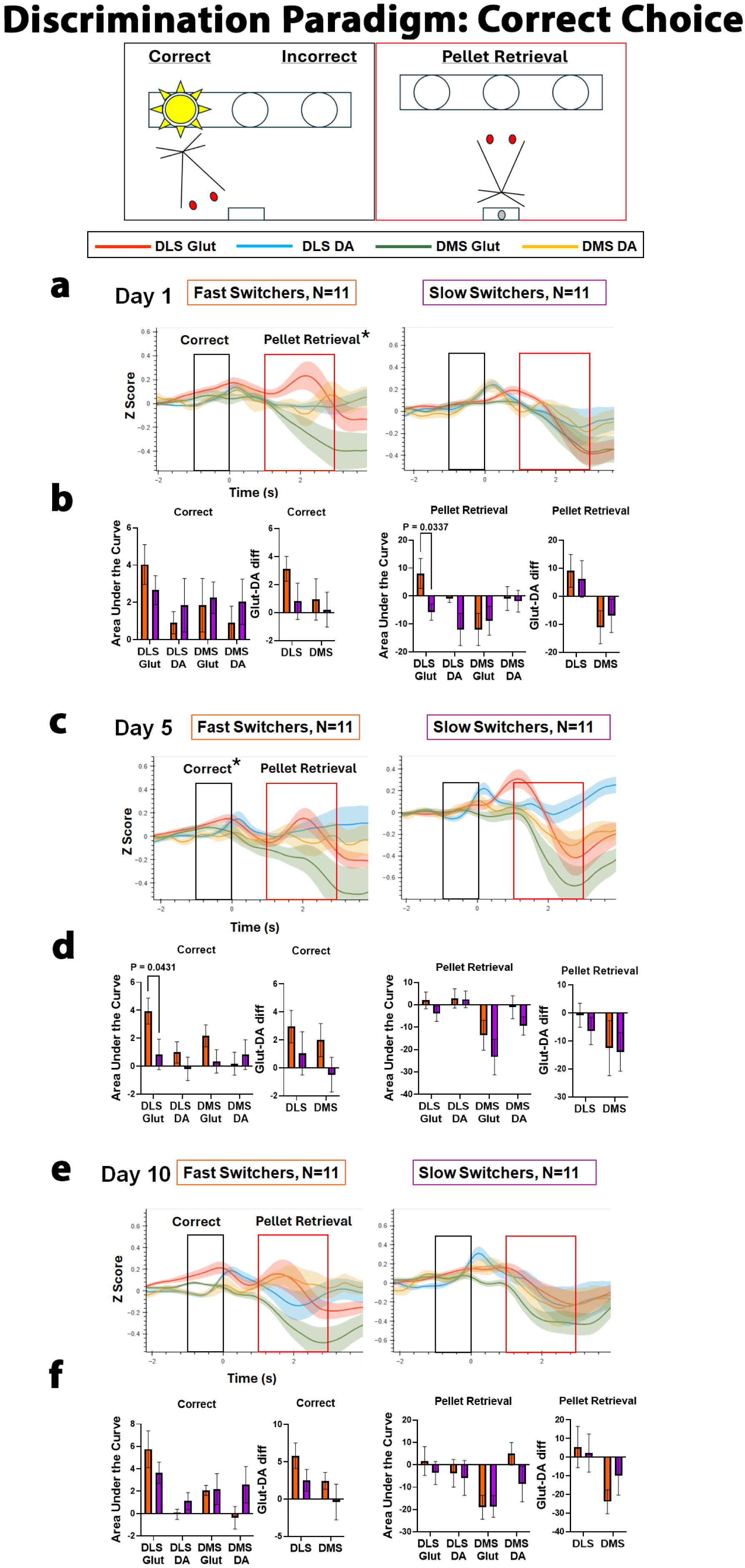
Fast switchers show elevated DLS Glut at correct choice and pellet retrieval: This figure depicts mean tracings just as in figure 2. **a,b)** Fast (orange) and slow (purple) switchers showed no differences in Glut or DA prior to a correct choice on day 1, but fast switchers had elevated DLS Glut release during pellet retrieval (8.1±5.4) compared to slow switchers (-5.8±2.9). **c,d)** On day 5, there was a significant difference in DLS Glut prior to a correct choice with an elevation in fast switchers (fast: 4.0±1.0) compared to slow switchers (slow: 0.8±1.1); however, there were no differences in the other release measures. **e,f)** By day 10 all measures normalized across rats.

### Strategy Switching Test: Fast and slow switchers differed in DLS Glut and DA release at trial start

We predicted that DMS and DLS DA and Glut signals would change as rats progressed through the strategy switching test, so we took 3 “snap shots” across the test. We separated signals from the first 30 trials (early), during ∼trials 60-90 (middle), and during the last 20 trials as the rats were reaching the performance criterion of 10 consecutive correct (late). To determine whether release measures when the rat initiated a trial during the strategy switching test would be different between rats that would go on to be fast and slow switchers, we compared DA and Glut release in the DMS and DLS across the early, middle, and late part of the strategy switching test. Rats that went on to be fast switchers began to show different Glut and DA dynamics in the DMS and DLS at trial start early in the test (Fig 4a-b), but this interaction reached significance during the middle portion of the test (F(3,54)=3.95, P=0.013, Fig. 4c). This effect was driven by a dynamic interaction between the release measures in the fast switchers, compared to a relatively quiescent response in the slow switchers. Comparing area under the curve (Fig. 4d), we found an elevation in DLS Glut release in the fast switchers (P=0.034), an elevation in DLS DA release in the slow switchers (P=0.004), and a greater difference in Glut relative to DA release in the DLS of fast switchers (P=0.012). The difference between the dynamic response in fast switchers and the quiescent response in the slow switchers continued late into the test (F(3, 54)=4.12, P=0.011; Fig. 4e). Again, this was driven by elevated DLS Glut (P=0.008) and DLS Glut-DA difference (P=0.013) in fast switchers and elevated DLS DA in slow switchers (P=0.037).

**Figure 4.**
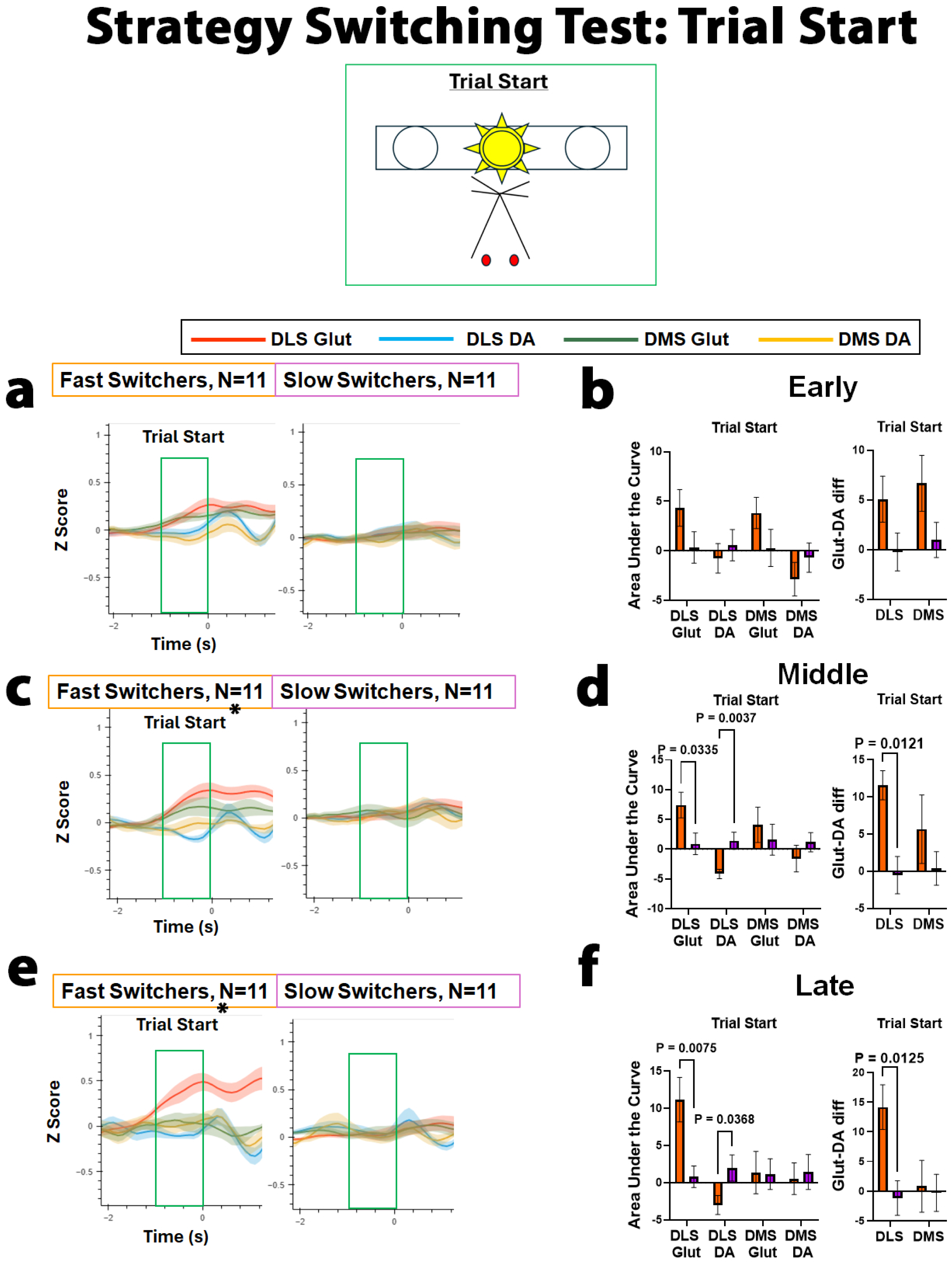
Fast switchers show elevated DLS Glut and a greater difference between Glut and DA at trial start during the strategy switching test: This figure depicts mean tracings aligned to trial start and is organized as prior figures. **a,b)** The differences in Glut and DA levels did not reach statistical significance at trial start during the early portion of the strategy switching test. **c,d)** In the middle portion of the test, DLS Glut was elevated in fast switchers (fast: 7.4±2.2, slow: 0.9±1.8) and DLS DA was elevated in slow switchers (fast:-4.2±0.8, slow: 1.4±1.4). There was also a significant difference when comparing Glut relative to DA in the DLS of fast switchers (11.6±2.0) that was not present in slow switchers (-0.5±2.5). There were no differences in the DMS. **e,f)** Late in the test, DLS Glut remained elevated in fast switchers both alone (DLS Glut - fast: 11.2±3.0, slow: 0.8±1.5) and relative to DLS DA (fast:-2.9±1.3, slow: 2.0±1.8).

### Strategy Switching Test: Fast switchers displayed test epoch-dependent changes in DA and Glut release prior to and after correct choices that were not seen in slow switchers

To determine whether specific neural signatures prior to choice and during pellet retrieval during the strategy switching test would be different between rats that would go on to be fast and slow switchers, we compared DA and Glut release in the DMS and DLS across the early, middle, and late part of the strategy switching test. Rats could make a correct choice in two ways; they could be correct when a response following the previous strategy happened to coincide with the new strategy or they could be correct following the new strategy. For example, if the light cue is in the left port when the rewarded strategy has switched from “the light is correct” to “left is correct,” the rat will be correct and receive a pellet reward even if they are still following the previous visual strategy (Fig 5a). If the rat chooses the left port when the light cue is on the right, then it is clear that they made a correct choice following the new response strategy. We hypothesized that DMS and DLS release measures could be different depending on whether the rat got a choice correct following the previous or new strategy, so we analyzed these correct choices and their subsequent pellet retrievals separately.

**Figure 5.**
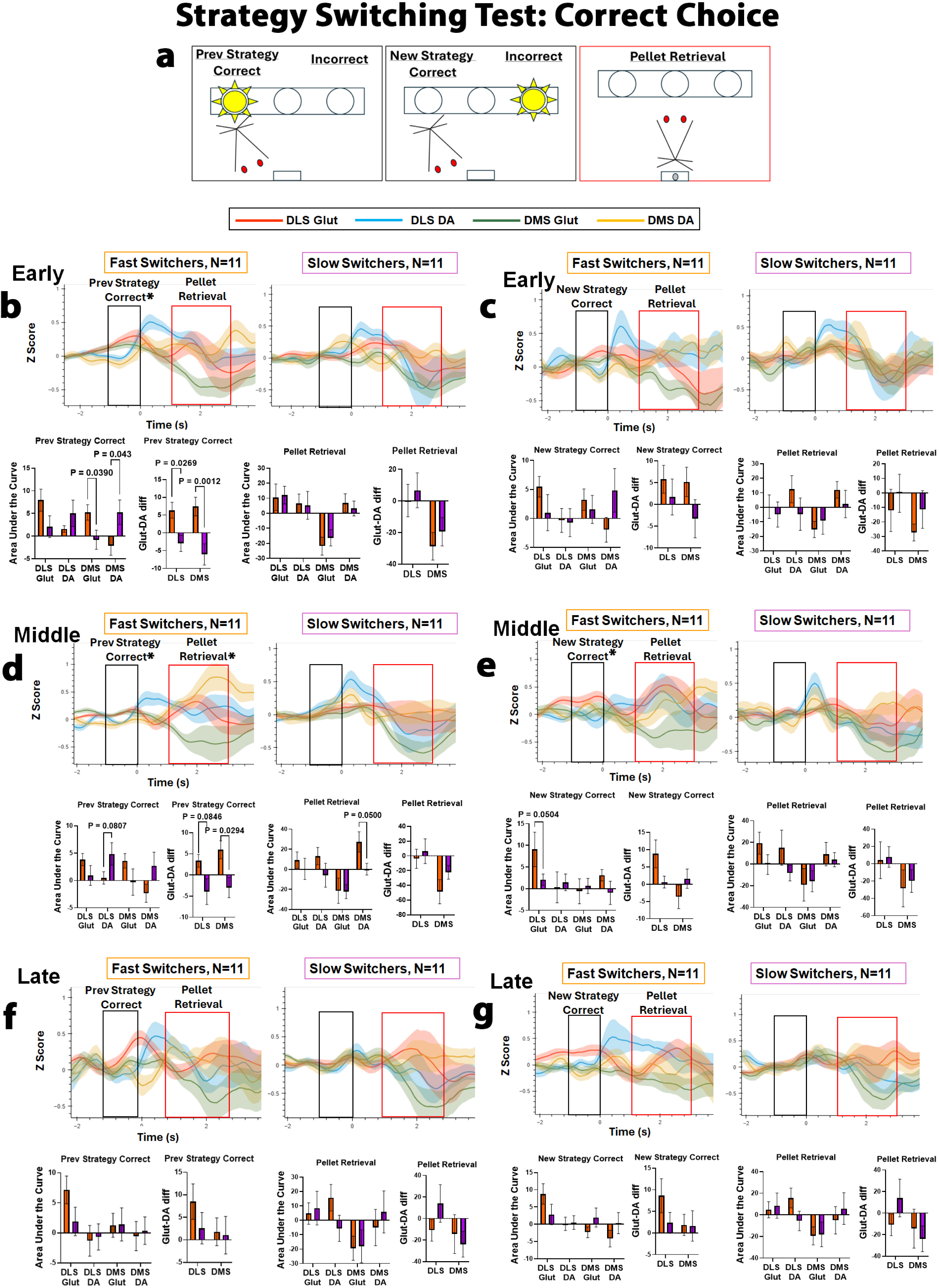
Fast switchers displayed test stage-dependent changes in DA and Glut release prior to and after correct choices that were not seen in slow switchers: **a)** This figure depicts mean tracings aligned to previous strategy and new strategy correct and is organized as prior figures. **b)** Early in the strategy switching test, fast switchers showed elevated DMS Glut (5.2±1.7) compared to slow switchers (-0.8±2.1), but reduced DMS DA release (-2.2±2.1) compared to slow switchers (5.2±2.7). DLS Glut (fast: 7.9±2.4, slow: 2.1±2.4) and DA (fast: 1.6±0.7, slow: 5.0±2.9) followed a similar pattern but did not reach significance. However, when quantifying Glut release, relative to DA release, between fast and slow switchers, we found that fast switchers had significantly higher Glut to DA ratios in both the DLS (6.4±2.2) and the DMS (7.4±2.6), while the slow switchers had elevated DA release, relative to Glut in both the DLS (-2.9±2.3) and DMS (-6.1±3.0). There were no differences in release measures during pellet retrieval during this epoch of the strategy switching test. **c)** Early in the strategy switching test, there were no significant differences when making a correct choice following the new strategy, although the patterns of release were very similar to those when making a correct choice following the previous strategy. **d)** In the middle portion of the strategy switching test, the fast switchers had decreased DLS DA release (0.5±1.1), while the slow switchers had elevated DLS DA release (4.9±2.1) prior to making a correct choice following the previous strategy. Fast switchers also featured elevated Glut release in DLS (fast: 3.9±1.1, slow: 0.9±1.8) and DMS (Glut-fast: 3.6±1.3, slow: - 0.3±2.4) and decreased DA release in DMS (fast:-2.3±1.7, slow: 2.7±2.5), but these effects did not reach statistical significance. Again however, fast switchers had elevated Glut in both the DMS and DLS when compared against DA release (DLS: 3.4±4.5; DMS: 5.9±6.4) compared to slow switchers (DLS:-3.9±10.2; DMS:-3.0±7.9). When looking at pellet retrieval, fast switchers had significantly elevated DMS DA release (27.4±10.1) compared to slow switchers (0.2±5.8). No other release measures within the pellet retrieval period were significantly different. **e)** In fast switchers, DLS Glut release was elevated prior to a correct choice following the new strategy (fast: 9.1±4.0, slow: 2.1±1.3), but there were no other significant differences. **f-g)** By late in the strategy switching test, dorsal striatal DA and Glut release measures were no longer significantly different in any measure.

*Early strategy switching test:* Rats that went on to be fast switchers had significantly different Glut/DA dynamics in DMS and DLS when they made a correct choice following the previous strategy (F(3,60)=4.72, P=0.005; Fig. 5b). Comparing area under the curve, we found an elevation in DMS Glut release in the fast switchers (P=0.039), and an elevation in DMS DA release in the slow switchers (P=0.043). Fast switchers also had a greater difference in Glut relative to DA release in both the DLS (P=0.027) and DMS (P=0.001). There were no differences during pellet retrieval and consumption during the early portion of the test. There were only a few instances of rats making a correct choice following the new strategy during the early portion of the test, as most rats continue to respond via their well-learned visual strategy. Thus, even though patterns of release were similarly different between fast and slow switchers, high variability and low power prevented statistically significant effects at “new strategy correct” (Fig. 5c).

*Middle strategy switching test:* Release measures during this portion of the test were similarly different between fast and slow switchers when they made a correct choice following the previous strategy (F(3,54= 4.020, P=0.012; Fig. 5d). Fast switchers featured a greater difference in Glut relative to DA release in both the DLS (P=0.085) and DMS (P=0.029), and slow switchers displayed elevated DA release in the DLS (P=0.081). Different from early in the test, however, fast switchers developed a large elevation in DMS DA release during Pellet Retrieval that was not seen in the slow switchers (P=0.05). The patterns in release measures were similar prior to making a correct choice following the new strategy (Fig 5d), with greater DLS Glut release prior to a correct choice (P=0.05) and a greater degree of activity during Pellet Retrieval in the fast switchers. Again, however, because there were fewer of these types of responses, there was less power to detect statistical significance.

*Late strategy switching test:* Different from the two earlier snapshots, these tracings are related to how the rat is performing in the test, rather than being aligned to a specific range of trials. Release measures during this portion of the test still appeared somewhat different; however, the dynamics of the slow switchers prior to a correct choice and during pellet retrieval became similar enough to the fast switchers that there were no longer any statistical differences. There were no differences prior to or following an incorrect choice (no pellet) across any time epoch (Fig.S3).

### DLS and DMS Glut release both diverge and converge across discrimination and strategy switching

Many studies over the years have highlighted a competitive balance between the DLS and DMS in habitual versus goal-directed behaviors, respectively. However, other studies have highlighted the possibility of more cooperative actions of these two dorsal striatum regions in action selection (reviewed in^23^). We analyzed DLS and DMS Glut release across time and we find evidence of both a divergence and convergence of Glut release across discrimination performance and strategy switching. During the discrimination paradigm and late in the strategy switching test in fast switchers, we see evidence of a competitive nature between the DLS and DMS, with DLS Glut release being significantly higher than DMS Glut at trial start (F(1,20)=6.40, P=0.0199) and correct choice (F(1,20)=15.3, P=0.0009) across days (Fig.S4a). During the early and middle portions of the strategy switching test, when uncertainty is the highest, we see a convergence of DMS and DLS Glut release, particularly prior to making a correct choice (F(8,150)=1.96, p=0.055; Fig.S4c).

### Specific differences in DLS and DMS Glut and DA release measures predict strategy switching performance and punishment resistant cocaine use

Using a correlation matrix, we found that the majority of the measured differences in DMS and DLS DA and Glut release between fast and slow switchers were significantly correlated with the number of strategy switching errors the rat would go on to commit, the number of punished cocaine infusions the rat would go on to take, or both (Fig. 6a). We used a Chi-Square test to identify which release measures were the strongest predictors of strategy switching performance and punishment sensitivity (Fig. 6b). The Chi-Square test corroborated the correlation matrix data, demonstrating that several measures were top predictors of both strategy switching and punishment sensitivity (Fig. 6c). To determine if we could achieve 100% predictability by combining neurotransmitter metrics, we created a linear discriminant analysis model. The model examined all possible combinations of the 14 metrics and used eight-fold cross-validation to evaluate performance of the classification models. We found that two different combinations of 2 metrics yielded 100% predictability for strategy switching performance (Fig. 6c-d) and two different combinations of 5 metrics yielded 100% predictability for punished cocaine use (Fig. 6e-f). Furthermore, hundreds of more combinations of 2-5 metrics were able to perfectly categorize strategy switching performance and categorize punishment sensitivity at better than 90% accuracy.

**Figure 6.**
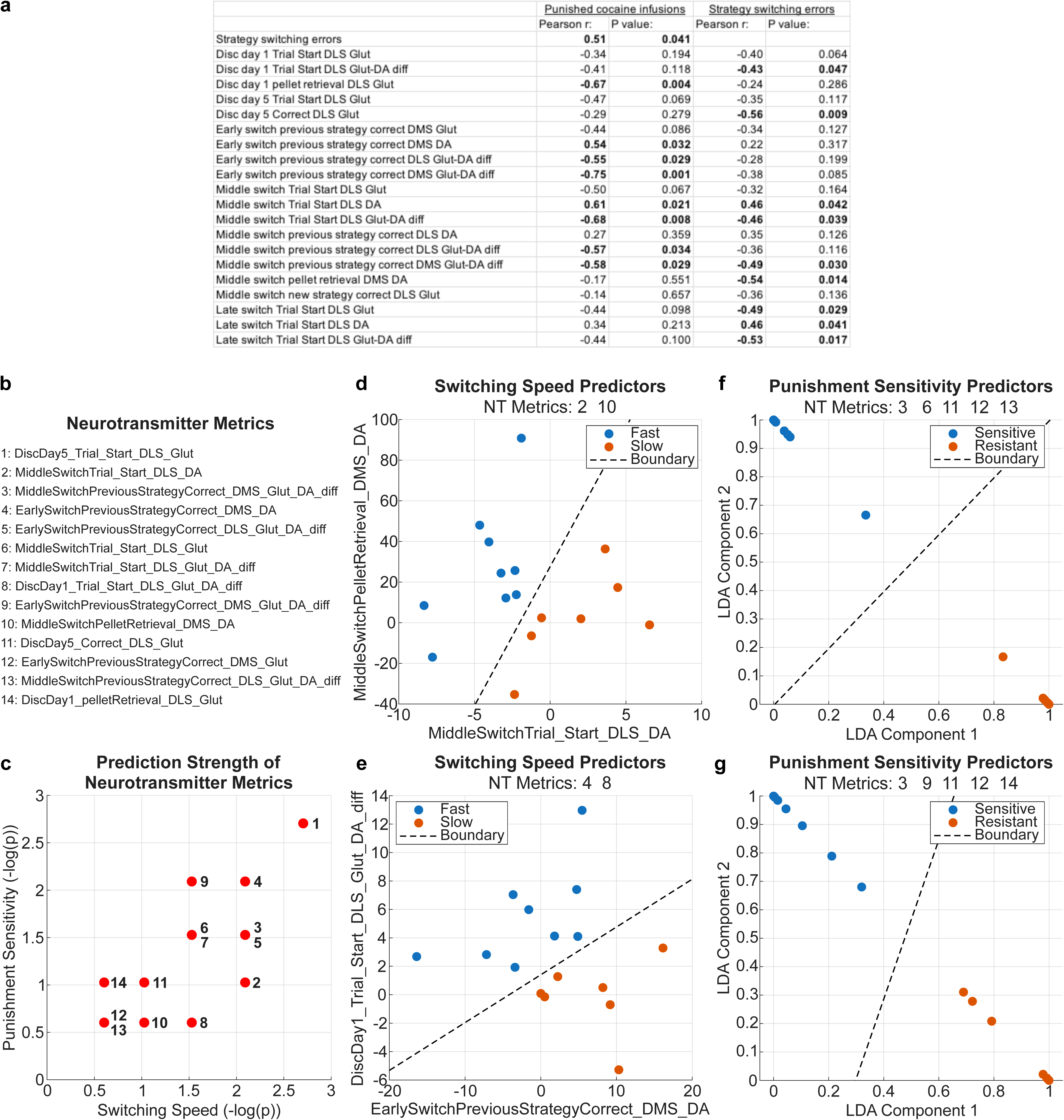
Specific combinations of DLS and DMS Glut and DA release measures predict strategy switching performance and punishment resistant cocaine use: **a)** Correlation matrix results with r and p values indicated. **b)** Fourteen neurotransmitter release metrics that were correlated with strategy switching performance and/or punishment sensitivity were tested for their prediction strength using a Chi-Square test. This list of metrics is the legend for the metric numbers across the rest of the figure. **c)** The prediction strength of each in relation to punishment sensitivity (y-axis) and strategy switching (x-axis) were plotted. **d)** Linear Discriminant Analysis (LDA) was used to examine every combination of the 14 neurotransmitter metrics and it identified 2 combinations of 2 metrics that correctly categorized the strategy switching performance of all rats, metrics 2 and 10 and **e)** metrics 4 and 8. **f)** The model identified 2 different combinations of 5 metrics that correctly categorized the punishment sensitivity of all rats, metrics 3,6,11,12, and 13 and **g)** metrics 3,9,11,12, and 14. These metrics were combined into LDA component 1 and 2 in order to display the separation of the 5 metrics in a 2-D plane.

### Orbital frontal and intralaminar thalamic projections to dorsal striatum are responsible for timestamp-specific Glut release and are necessary for fast strategy switching and sensitivity to cocaine punishment

To determine the source(s) of the Glut measured at specific behavioral timestamps and determine whether they are necessary for fast switching and punishment sensitivity, we inhibited two Glut inputs to the DMS and DLS with pathway-specific DREADDs and measured the effects on the photometry signal, discrimination performance, strategy switching, and punished cocaine use. We chose to inhibit the ventral orbital frontal cortex (OFC) and intralaminar thalamus (IN Thal) projections to dorsal striatum because these regions send dense glutamatergic projections to the dorsal striatum that are parallel to the DMS and DLS^24–26^. They are also both known to be involved in some version of attention or behavior switching^13,27–35^. We infused a Cre-dependent, Gi-coupled DREADD into the OFC, a Cre-dependent KOR DREADD into the IN Thal, and a retrograde CRE virus into the DMS and DLS (Fig. 7a-b). Thus, injection of CNO, the designer drug for the Gi-coupled DREADD, would result in the inhibition of OFC inputs to dorsal striatum, and injection of SalB, the designer drug for the KOR DREADD, would inhibit IN Thal inputs to dorsal striatum. The same photometry viruses and fibers as used in the first experiment were also infused and implanted into the DMS and DLS, allowing the measurement of Glut and DA in response to inhibition of OFC and/or IN Thal inputs. Rats then underwent the discrimination paradigm and strategy switching test as in the first experiment, and we performed photometry recordings as above. Our classification model provided hundreds of different combinations of neurotransmitter metrics that perfectly categorized the strategy switching performance of every rat from the first experiment. We selected three 3-metric models that combined metrics that were taken from these rats when they were unmanipulated (vehicle conditions): discrimination day 1, day 5, and early in the strategy switching test (see Fig.S5). If 2 out of 3 models identified the rat as a fast switcher then we categorized them as a “putative” fast switcher, and vice versa. The model identified 7 putative fast (3 male/ 4 females) and 7 putative slow (5 male/ 2 female) switchers (Fig.S5). We analyzed the putative fast and slow switchers separately as we did in the first experiment.

**Figure 7.**
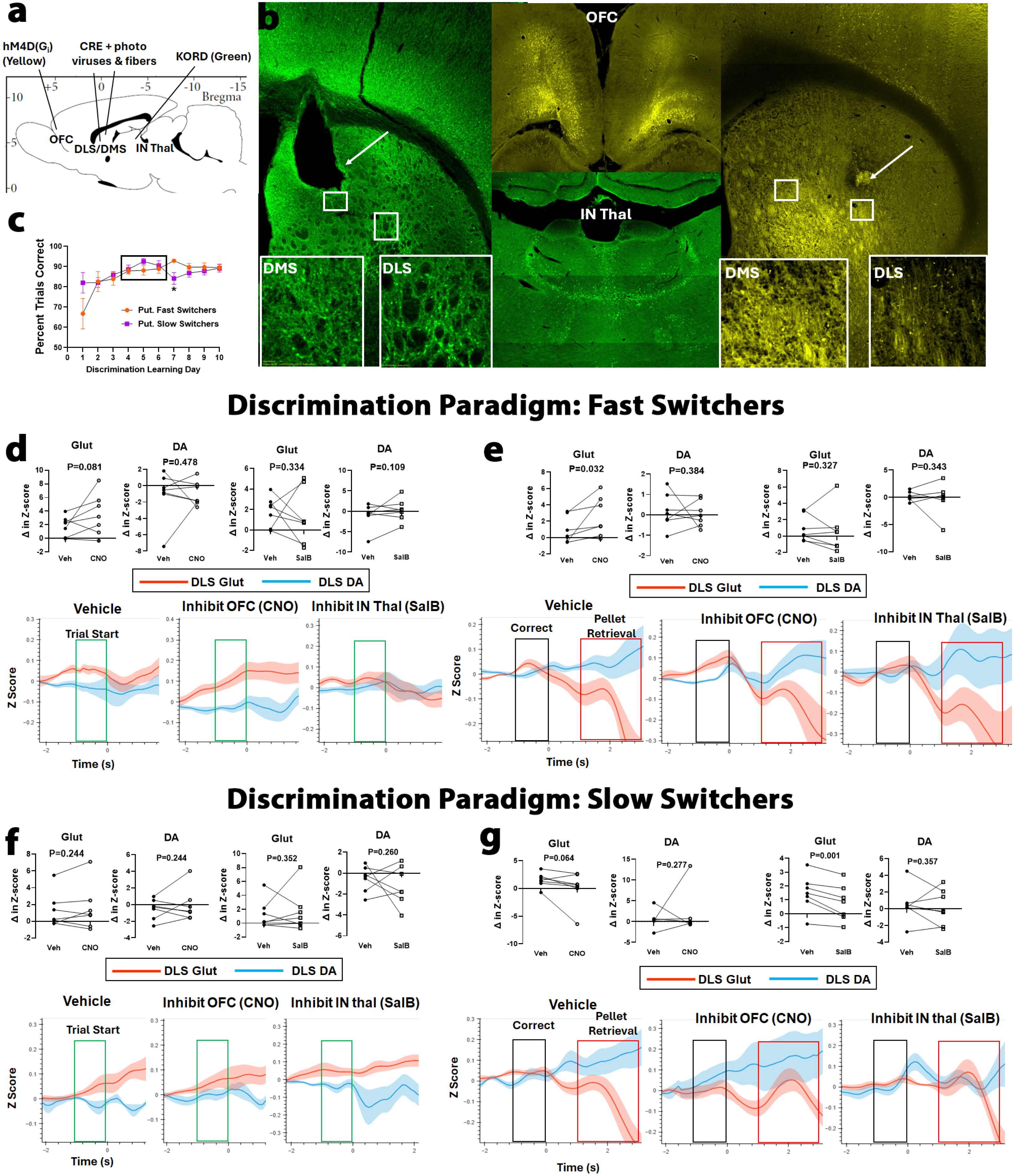
Inhibition of OFC and IN Thal inputs produce opposite effects on Glut release in DLS in putative fast and slow switchers: **a)** Virus/fiber infusion/implant map. **b)** Representative images showing green staining from IN Thal and yellow staining from OFC into both the DMS and DLS (insets). Arrows indicate fiber placements in the DMS and DLS. **c)** Inhibition of OFC and IN Thal inputs impaired slow switchers on day 7 (fast-92.9, slow-84.1; p=0.022). Mean tracings are aligned to trial start (green box), correct choices (black box), and subsequent pellet retrieval (red box) of DLS Glut (red) and DLS DA (blue). Data across rows are within subjects and represents the change in signal (Z-score) from no manipulation (vehicle, left) to inhibition of OFC (CNO, middle) and then inhibition of IN Thal (SalB, right) inputs to dorsal striatum. The tracings are quantified using area under the curve (Mean ± SEM) for the 1 or 2 seconds in the colored box, as indicated. Inhibition of OFC inputs increased DLS Glut in putative fast switchers (N=7) at both **d)** trial start (Veh: 1.84±0.55, CNO: 3.51±1.14) and **e)** correct choices (Veh: 0.89±0.61, CNO: 2.51±0.91) but did not affect DA release. Inhibition of IN Thal inputs had no effect on Glut release or DA release. **f)** Inhibition of OFC (Veh: 1.49±0.49, CNO: 0.08±1.15) and IN Thal (SalB: 0.78±0.49) inputs reduced glut release at correct choice in putative slow switchers (N=7) but had no effect on DA release. **g)**There was no effect on Glut at trial start.

*Discrimination paradigm:* We found the greatest difference between eventual fast and slow switchers on discrimination day 5, so between days 4-6 rats underwent 1 day with a vehicle injection, one day with a CNO injection, and one day with a SalB injection (order counterbalanced across rats) to compare the effects on Glut release both within and across rats. Discrimination performance was impaired in the putative slow switchers, such that there was a significant reduction in the number of correct responses on the day after the CNO/SalB injection days (F(9,117)=2.531, p=0.011; Fig. 7c). Learning in the putative slow switchers was reset to some degree as their mean percent trials correct dropped from 91% on day 6 to 84% on day 7, similar to what it had been on days 1 and 2 (82%). Discrimination performance was not affected in the putative fast switchers. In accordance with the behavioral data, inhibition of OFC inputs actually increased Glut release in the DLS of putative fast switchers at both trial start (t(6)=1.59, p=0.081; Fig. 7d) and prior to a correct choice (t(6)=2.28, p=0.032; Fig. 7e). In putative slow switchers, inhibiting OFC (t(6)=1.76, p=0.064) and IN Thal (t(6)=4.84, p=0.001) reduced glutamate release in DLS prior to a correct choice (Fig. 7g). The only similarity between putative fast and slow switchers was that inhibition of IN Thal inputs reduced glutamate release in DMS prior to a correct choice in both fast (t(6)=3.28, p=0.008) and slow switchers (t(6)=2.23, p=0.034; Fig.S6).

*Strategy switching test:* We found the greatest difference between eventual fast and slow switchers during the early (trials∼1-30) and middle (trials∼60-90) portions of the test. So, we split the first 90 trials of the strategy switching test into 3 sets (1-30, 31-60, 61-90) and each rat underwent 1 set with a vehicle injection, 1 set with a CNO injection, and 1 set with a SalB injection (order counterbalanced) to compare the effects on Glut release both within and across rats. Behavior was stopped for the day after 30-trial sets with injections of CNO or SalB to allow drug washout before the next set of 30 trials on the following day. In putative fast switchers, inhibition of both IN Thal (t(5)=3.05, p=0.014) and OFC (t(4)=2.79, p=0.025) inputs reduced Glut release in the DLS at trial start (Fig. 8a) and prior to a correct choice following the previous strategy (OFC:t(4)=2.25, p=0.044; IN Thal:t(5)=3.02, p=0.015; Fig. 8b). The effects in DMS were more specific, with IN Thal inhibition reducing Glut release at trial start (t(6)=1.85, p=0.057; Fig. 8c) and OFC inhibition reducing Glut release prior to a correct choice following the previous strategy (t(4)=6.23, p=0.002; Fig. 8d). Interestingly, we also measured several changes in DA release in DMS and DLS following inhibition of OFC and IN Thal inputs. The most consistent were an increase in DMS DA at trial start (t(4)=4.20, p=0.007; Fig. 8c) and a decrease in DMS DA during pellet retrieval (t(4)=2.95, p=0.021; Fig. 8d), both following inhibition of OFC inputs to DMS. Neither inhibition of OFC or IN Thal inputs had any consistent effect on the Glut measured prior to a correct choice following the new strategy, although fewer rats sampled the new strategy during this portion of the test (Fig.S7). There were, however, reductions in DMS DA during pellet retrieval (Fig.S7b). Neither inhibition of the OFC or IN Thal inputs had any consistent effect on Glut release at any behavioral timestamp in the putative slow switchers (Fig.S8), which was expected given the general lack of Glut release in slow switchers. After the first 90 trials, all rats finished the strategy switching test without further manipulation. Putative fast switchers were impaired in their strategy switching performance relative to the fast switchers from the first experiment (P=0.0067), whereas putative slow switchers were improved (P=0.0009; Fig. 8e).

**Figure 8.**
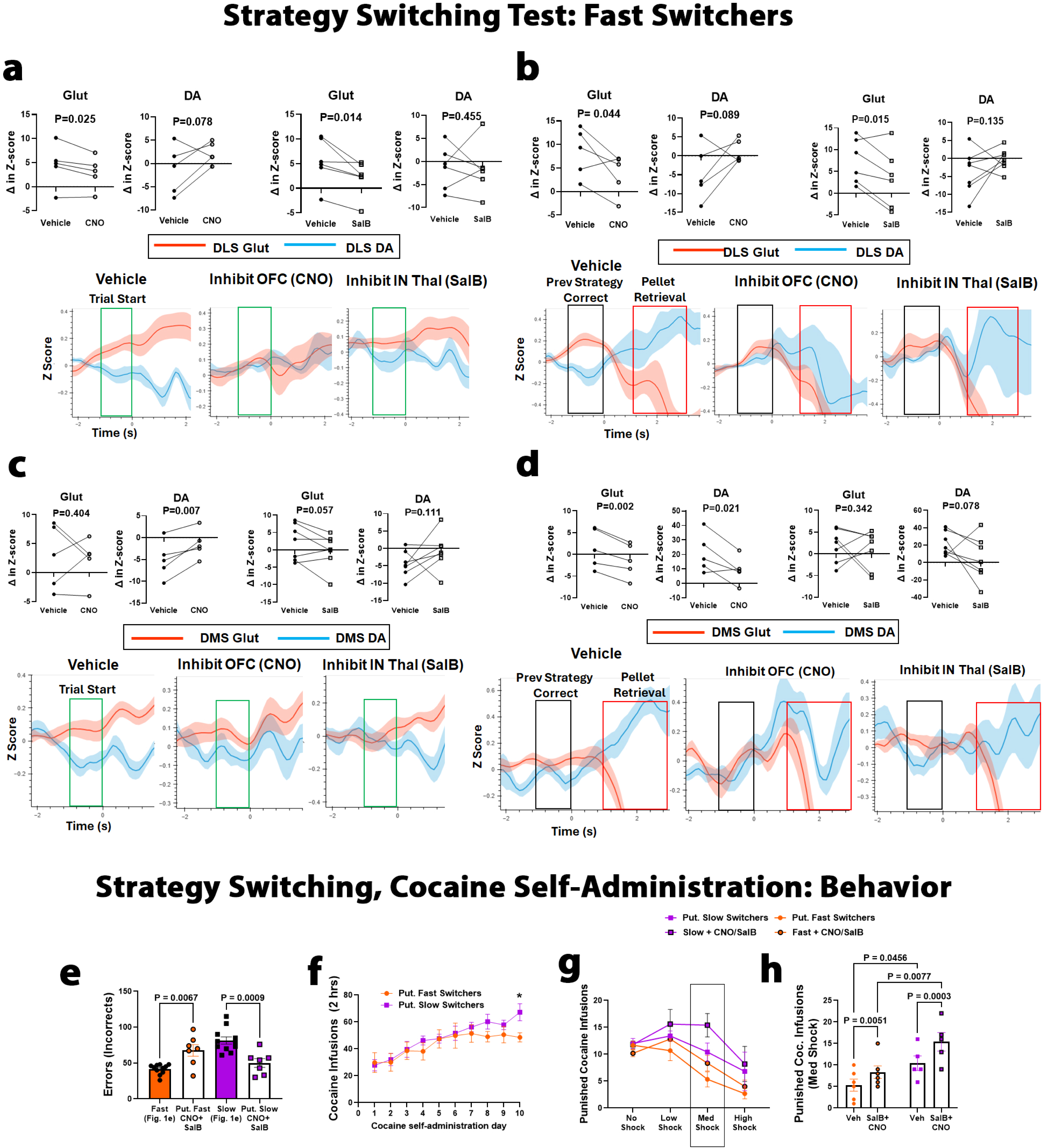
OFC and IN Thal projections to dorsal striatum are responsible for behavioral timestamp-specific Glut release in putative fast switchers and are necessary for fast strategy switching and sensitivity to cocaine punishment: This figure is organized as figure 7. **a)** Inhibition of OFC inputs decreased **<u>DLS</u>**Glut (Veh: 4.45±1.99, CNO: 2.93±1.51) and increased DA (Veh:-1.36±2.36, CNO: 2.26±1.02) at trial initiation, whereas inhibition of IN Thal inputs decreased DLS Glut (Veh: 5.46±1.91, SalB: 2.12±1.46), but did not affect DA. **b)** Inhibition of OFC inputs (Glut-Veh: 8.36±2.30, CNO: 3.69±1.93) and IN Thal inputs (Veh: 7.42±2.10, SalB: 3.48±2.77) also decreased **<u>DLS</u>** Glut at correct choices following the previous strategy, but only OFC inhibition increased DLS DA (Veh:-4.46±3.25, CNO: 1.24±1.40). **c)** Inhibition of OFC inputs did not reduce **<u>DMS</u>** Glut at trial initiation (Veh: 2.75±2.48, CNO: 2.19±1.70) but did result in an increase DA release (Veh:-5.08±1.89, CNO:-1.45±1.44). Inhibition of IN Thal inputs decreased DMS Glut at trial initiation (Veh: 2.31±1.96, SalB:-0.26±1.86) but did not significantly affect DA release. **d)** Inhibition of OFC inputs decreased DMS Glut prior to a correct choice following the previous strategy (Veh: 1.40±2.03, CNO: - 1.36±1.74) and decreased <u>DMS DA during the pellet retrieval</u> period (Veh: 20.88±5.96, CNO: 9.25±4.15). DA release at previous strategy correct was not affected by OFC inhibition. Inhibition of IN Thal inputs did not significantly reduce DMS Glut prior to a correct choice following the previous strategy but did still reduce <u>DMS DA during the</u> <u>pellet retrieval</u> period (Veh: 21.79±5.10, SalB: 4.27±9.70). DA release at previous strategy correct was not affected by IN Thal inhibition. **e)** Putative fast switchers that finished the strategy switching test after treatment with CNO and SalB (errors: 67.8±7.96) were significantly impaired compared to fast switchers from the first experiment (orange bar: 41.6±2.1). Putative slow switchers (50.0±6.03) were improved relative to slow switchers from the first experiment (purple bar, 81.4±5.0). **f)** Putative fast and slow switchers took similar numbers of cocaine infusions across the 10 self-administration days; however, slow switchers (67.2±6.31) took more infusion on day 10 than fast switchers (48.57±3.41; p=0.039). **g)** Putative slow switchers took more punished cocaine infusions overall. **h)** Due to the short half-life of SalB, the effect of inhibiting both IN thal and OFC inputs to dorsal striatum was limited to the medium shock bin (30 minutes total). Inhibition of OFC and IN Thal inputs increased the number of punished cocaine infusions that putative fast (P=0.005) and slow switchers (P=0.0003) relative to their to vehicle injection. Slow switchers took more punished cocaine infusions than fast switchers under both vehicle (P=0.046) and SalB+CNO (P=0.008) conditions.

*Cocaine Self-administration:* Following the strategy switching test, all rats were implanted with jugular vein catheters and trained to self-administer cocaine as before. After acquisition, all rats were given continuous access to cocaine for 2 hours across 10 days, as this revealed the largest difference between fast and slow switchers in the prior experiment. Putative slow switchers took similar numbers of cocaine infusions as fast switchers across the first 7 days. Slow switchers took more cocaine infusions across the last 3 days, which reached significance on day 10 (P=0.039; Fig. 8f). Rats then underwent a first punishment resistance test, followed by 4 further days of self-administration and then a second punishment resistance test. Each rat underwent 1 punishment resistance test following a vehicle injection and one test following an injection of both CNO and SalB (order counterbalanced). Putative slow switchers took more punished cocaine infusions overall (F(1,9)=8.94, p=0.015), as in the first experiment; however, injection of CNO and SalB increased the number of punished cocaine infusions taken in both putative fast and slow switchers (Fig. 8g). CNO injection was given prior to the start of the punishment resistance test, but, because of SalB’s short half-life, it was given just prior to the medium shock intensities. Thus, we can observe an effect of inhibiting OFC inputs across the whole test but can only compare the effect of inhibiting both OFC and IN Thal projections during the medium shock intensities. Treatment with CNO and SalB increased the number of punished cocaine infusions across all rats (F(1,9)=43.64, p<0.0001) and within both putative fast (p=0.005) and slow (p=0.0003) switchers (Fig. 8h).

## Discussion

We showed that rats that were inherently slow at strategy switching were also the most likely to develop punishment resistant cocaine use. Photometry recordings during discrimination and the strategy switching test showed that fast switchers had elevated dorsal striatal Glut release and reduced DA release at trial initiation (DLS) and prior to a correct choice (DMS and DLS). They also featured dynamic fluctuations in release measures to the positive feedback of receiving the pellet reward, particularly an increase in DA release in the DMS. Slow switchers showed the opposite pattern, with elevated DMS and DLS DA release prior to a correct choice and a quiescent response at trial initiation and to positive feedback. Many of these release measures were significant predictors of either strategy switching performance, punishment resistant cocaine use, or both as indicated by our correlation matrices, Chi-Square test, and classification models. Interestingly, quantifying the dynamic relationship of Glut release relative DA release was often a better individual predictor than measuring either DA or Glut individually. Furthermore, combining 2-5 release metrics allowed our classification model to categorize switching speed and punished cocaine use with 100% accuracy. This emphasizes the importance of measuring the dynamic relationship between Glut and DA release across several behavioral measures. In a separate experiment, we showed that rats categorized by our model as fast and slow switchers showed differential recruitment of Glut inputs to the dorsal striatum during discrimination and strategy switching. Putative fast switchers showed significant contributions from both the OFC and the IN Thal in DLS and DMS across several strategy switching timestamps but showed little to no recruitment of these regions during performance of the discrimination paradigm. Putative slow switchers showed little to no contribution from the OFC and IN Thal during the strategy switching test but showed significant contribution of both regions to the DLS during the discrimination paradigm. There was also a differential effect on behavior across these rats that aligned with the reductions in Glut release, with OFC and IN Thal inhibition impairing putative fast switchers on the strategy switching test only and putative slow switchers on the discrimination paradigm only. Interestingly, inhibition of OFC and IN Thal inputs to dorsal striatum increased the number of punished cocaine infusions taken across all rats. This emphasizes our correlation matrix and modeling data showing that Glut/DA interactions during both the discrimination paradigm and strategy switching can predict punished cocaine taking but also demonstrates that the source of the Glut across discrimination performance and strategy switching is likely different. Based on these data, we propose a model where the combination of projection-specific Glut release and a DA dip at behavioral initiation (DLS) or at a decision point (DMS and DLS), with a subsequent increase in DA and Glut during the feedback period, represents a neurochemical “signature” of flexibility that predicts a greater ability to adapt to changing contingencies (e.g. a strategy switch or punishment addition, Fig. 9). In contrast, dorsal striatal DA release leading to a decision point, with a quiescent response at behavioral initiation and during the feedback period, represents an inflexible signature (Fig. 9).

### Glut-DA dynamics for flexible behavior

Our data suggest that elevated Glut release, with a concomitant reduction in DA release, at trial initiation or prior to choice predicts both fast strategy switching and punishment sensitivity, whereas elevated DA release at these timepoints predicts the opposite. This suggests that the timing of the interaction of DA and Glut release at these timepoints is critical. We have two suggestions for why this might be. First, an increase in Glut paired with a decrease in DA could prime the striatum to maximally attend to trial-by-trial feedback. The striatum is primarily made up of medium spiny projection neurons (MSN) that have a hyperpolarized resting potential^36^. Prior studies have shown that Glut inputs to striatum can reduce this resting potential, putting them in an “upstate” and making it easier for them to fire^37^. A dip in DA levels can further favor Glut release from these inputs by releasing them from presynaptic D2-mediated inhibition^38,39^, which specifically favors subsequent input from corticostriatal projections^40^. Because of this, feedback-related elevations in phasic Glut and DA release can facilitate LTP and learning^41,42^. A quiescent Glut signal with small increases in DA at trial start or prior to a choice would instead result in further inhibition of Glut release via presynaptic D2’s^38,39^, leaving MSN’s in a hyperpolarized “downstate”. We see evidence of this notion in our own data. For example, fast switchers, which feature the increase in Glut and decrease in DA at trial start and prior to a correct choice, also feature dynamic fluctuations in Glut and DA release during pellet retrieval, including a significant elevation in DMS DA. Furthermore, slow switchers, which feature elevated DA and no increase in Glut at trial start and prior to a correct choice, also feature no feedback-related activity during pellet retrieval. Lastly, when the Glut inputs from OFC and IN Thal were removed from putative fast switchers, we showed a decrease in Glut release at trial start and prior to choice which coincided with a reduction in feedback-related DMS DA release at pellet retrieval. This occurred without manipulating DA inputs directly, suggesting that it was a result of the Glut inhibition (See Fig. 8d, S7b).

A second possibility is that the increase in Glut and decrease in DA could facilitate the indirect pathway, leading to a greater inhibition of the previously learned strategy. MSN’s are categorized by the presence of D1 or D2 receptors. D1 receptor-containing MSN’s facilitate movement and action selection via activation of the direct pathway and D2 receptor-containing MSN’s inhibit movement and refine action selection via activation of the indirect pathway^43,44^. A dip in DA release has been shown to facilitate LTP in D2 MSN’s and LTD in D1 MSN’s^45^, thus favoring the indirect pathway and motor inhibition. Furthermore, thalamic inputs to the dorsal striatum synapse onto both MSNs and cholinergic interneurons (CIN’s)^46^. CIN activation favors corticostriatal signaling, ultimately leading to activation of the indirect over the direct pathway^46^. This has been suggested by others as a mechanism for salient information to lead to action suppression to facilitate shifts in attention and behavior^46,47^. Again, there is evidence of this notion in our data. We showed that the Glut we measured at trial start and prior to a correct choice following the previous strategy was from cortex (OFC) and thalamus (IN Thal), both of which would favor activation of the indirect pathway. Importantly, these inputs were not responsible for the Glut measured when fast switchers were making a correct choice following the new strategy (Fig.S6). Thus, the increase in thalamocortical Glut inputs with a DA dip in the fast switchers could lead to activation of CINs and D2 MSNs, potentially improving the fast switcher’s ability to inhibit responding via the previous, well-learned strategy despite getting that trial correct. The slow switchers, which did not feature Glut increases from thalamus or cortex during the strategy switching test (Fig.S8), had increased DA release prior to choice. This would be expected to activate D1 MSN’s leading to direct pathway activation^43,44^, inhibition of corticostriatal input^40^, and perseveration on the previous strategy^46^.

### Differential effect of OFC and IN Thal inhibition on Glut release in putative fast and slow switchers

Inhibition of OFC inputs resulted in diminished Glut release in both the DMS and DLS prior to a correct choice following the previous strategy, as well as at trial start in the DLS. Inhibition of the IN Thal resulted in diminished Glut release at trial start in both the DMS and DLS, as well as prior to a correct choice following the previous strategy in the DLS. However, this was only the case in putative fast switchers. Inhibition of the OFC and IN Thal had little to no effect on Glut release during the strategy switching test in putative slow switchers. Slow switchers showed significantly less Glut released during the strategy switching test timestamps in the first experiment, suggesting that Glut levels may not have been able to be reduced due to a floor effect. That said, rats in the putative fast switcher group had a range of Glut release magnitudes, including some that were quite low, which were reduced by chemogenetic inhibition of the OFC and/or IN Thal inputs. Furthermore, in putative slow switchers, inhibition of OFC and IN Thal inputs reduced DLS Glut during the discrimination paradigm, and this did not occur in the putative fast switchers. In fact, inhibition of OFC inputs actually increased Glut release in the DLS at trial start and prior to a correct choice in fast switchers. This suggests that there is a fundamental difference between the dorsal striatum input architecture of fast and slow switchers, with opposite activation patterns of OFC and IN Thal inputs during discrimination and strategy switching in these two rat phenotypes. We know that the OFC tracks changes in action value and is necessary for shifts in behavioral strategies^29^ and probabilistic reversal learning^28^. The OFC also supports state representations as a means to promote new learning when outcomes diverge from expectations^48^, as would happen when contingencies change during strategy switching. The IN Thal is necessary for initiation^13^ and execution^14^ of well-learned behaviors, redirecting attention^31,32^ and integrating new and old information^27^ in response to contingency shifts. Thus, a contribution of these regions during the strategy switching test in fast, but not slow switchers is likely at the heart of the difference between “flexible” and “inflexible” rats.

### Differential effect of OFC and IN Thal inhibition on behavior in putative fast and slow switchers

A curious finding was that rats predicted by our model to be slow switchers were better at the strategy switching test than the slow switchers from our first experiment. Is it possible that our model was incorrect in its categorization of these rats or could there be another explanation for this counterintuitive phenomenon? We used 3 different 3-metric models, which perfectly categorized the rats from the first experiment, to categorize these rats to minimize this risk. Furthermore, 10 of the 14 rats were categorized as fast or slow by all 3 models, suggesting a high level of confidence in the classifier of at least those 10. Furthermore, the mean tracings for the putative fast and slow switchers look remarkably similar to the respective fast and slow switchers from the first experiment. The putative fast and slow switchers also demonstrated interesting differences in the contributions of OFC and IN Thal inputs during the discrimination paradigm and strategy switching test outlined above. These effects were consistent across rats within each group, suggesting a difference in input architecture and utilization that aligns with their behavior. Thus, although it is still possible, we do not believe that the putative fast and slow switchers were miscategorized. Instead, we believe the improved strategy switching in the putative slow switchers may be because their performance in the discrimination paradigm was destabilized by the chemogenetic inhibition. We found that inhibition of both OFC and IN Thal inputs during the discrimination paradigm reduced Glut release in the DLS of putative slow switchers. This led to a dip in performance from day 6 (91%), such they got a similar percentage of trials correct on day 7 (84%) as they did on days 1 and 2 (82%). Given that activity in the DLS increases across learning^17^, at the initiation of a well-learned behavior^15^, and across performance of the discrimination paradigm (Fig.S4), and that DLS inhibition increases rats’ sensitivity to changes in action-outcome contingency^49^, it may be that the reduction in DLS Glut could have reset how “well-learned” the visual strategy was, increasing the putative slow switchers’ sensitivity to the strategy switch. Importantly, the putative fast switchers did not show a reduction in DLS Glut at any timestamp during the discrimination paradigm, nor any dip in performance on any discrimination day.

### Dual effect of DMS DA release

DA release in the striatum is necessary for both learning and cognitive flexibility by strengthening the association of actions with rewarding outcomes^50^, which is separate from movement-related signaling^51^. We see evidence of this as well, as the rats that showed the greatest elevations in DMS DA release during pellet retrieval switched strategies the fastest (Fig. 6), and the reductions in strategy switching performance after OFC or IN Thal inhibition were accompanied by significant reductions in DMS DA release during pellet retrieval (Fig.8d, S7b). The more paradoxical finding may be that DMS DA released at trial start or prior to choice predicted both slow strategy switching and punishment resistance. This finding may seem to be at odds with prominent theories in the literature; however, two recent studies lend credence to the idea that DMS DA release can predict inflexibility^52^ and resistance to punishment^53^. For example, one study showed that the combination of elevated DMS DA release to a distal seeking cue (seeking lever extension) and decreased DMS DA levels at pellet delivery significantly predicted which rats would show habitual behavior during a habit test^52^. The opposite pattern predicted which rats would be goal-directed, and these patterns increased in magnitude across training^52^. Furthermore, optogenetic stimulation of DMS DA release during the distal cue seeking accelerated habit formation^52^. Moreover, another found that DMS DA release during a rewarded nosepoke (RI30 schedule) early in training predicted which rats would be punishment resistant on a first probe test^53^, and that optogenetic stimulation of DMS DA release accelerated the development of punishment resistant sucrose taking^53^. Interestingly, all mice showed an increase in DMS DA release by the end of training, indicating that the timing of the DMS DA release, i.e. when in training it developed, was critical for whether a mouse would be resistant or sensitive to punishment^53^. Thus, in both cases and similar to our results, the timing of the DMS DA release predicts flexibility. When it occurs early in training^53^, early in a behavioral progression^52^, or prior to a choice (Fig.5b), it is indicative of an inflexible, punishment resistant phenotype (Fig.6). When it develops late in training^53^ and occurs to provide feedback about an action^50–52^ (Fig. 5d), it is indicative of sound learning and a flexible, goal-directed phenotype (Fig.6).

### Summary

Across learning and strategy switching, elevations in Glut, with reductions in DA, at trial initiation (DLS) and prior to choice (DMS and DLS), with dynamic fluctuations in DMS DA and DLS Glut to positive reinforcement, represented a “signature of flexibility” that predicted both fast strategy switching and cocaine seeking that was sensitive to punishment. Elevations in DLS and DMS DA at these respective points, with a quiescent response to feedback, represented a “signature of inflexibility” that predicted slow switching and resistance to punishment. The differences in neurochemical signatures were accompanied by differences in input architecture and utilization across discrimination and strategy switching and represent inherent differences between a group of unmanipulated male and female rats. As such, these data describe a pattern that could be measured and/or manipulated to predict and treat SUDs in humans.

## Acknowledgements

This work was funded by NIH grants P50DA046346 and R01DA058955.

## Methods

### Subjects

Adult (>PND70) male and female Sprague-Dawley rats (Envigo, Frederick, MD) were used for all experiments. Confidence intervals were used to compare between sexes (see results). Both sexes were determined to be part of the same interval, as we have shown previously^54,55^, and were combined. We also did not observe sex differences in cocaine-motivated behaviors under the experimental conditions we used, similar to our prior work^28^. All rats were pair-housed unless otherwise noted and were maintained on a 12-12hr light-dark cycle, with unlimited access to food chow and water, except during behavioral testing, as described below. Experiments were performed during the dark cycle. Experimental procedures were approved by the Institutional Animal Care and Use Committee of the University of Pittsburgh according to National Institute of Health Guide for the Care and Use of Laboratory Animals.

### Virus infusion surgery

All viral constructs were purchased from Addgene. Prior to any behavioral training, Rats underwent an initial virus infusion surgery using our standard protocols^18,56–58^ where they received infusions of viral vectors that contain photo-emitting biosensors to detect DA (red-AAV_9_-hsyn-GRAB_rDA1m, 140556-AAV9; 0.5 µL at titer ≥ 1×10¹³ vg/mL, deposited by Yulong Li)^59^ and Glut (green-AAV_9_-hSynapsin-SF-igluSnFR.A184S, 106174-AAV9; 0.5 µL at titer ≥ 1×10¹³ vg/mL, deposited by Loren Looger)^60^ into the DMS of one hemisphere (AP:-0.2, ML: ±2.2, DV:-4.5) and the DLS of the other hemisphere (AP: +0.8, ML: ±2.8, DV:-5.0). In the same surgery, optic fibers (200µm diameter 0.37 NA, Doric) were implanted into the DMS and DLS (one fiber/ region, Fig. 1a).

For the final experiment only, rats were also infused with AAV_8_-hSyn-dF-HA-KORD-IRES-mCitrine (65417-AAV8; titer ≥ 7×10¹² vg/mL, deposited by Bryan Roth)^61^ into the Intralaminar thalamus (IN thal, AP:-4.1, ML: ±1.2, DV:-6.2; 0.5 µL/ hemisphere) and AAV_8_-hSyn-DIO-hM4D(Gi)-mCherry (44362-AAV8; titer ≥ 1×10¹³ vg/mL, deposited by Bryan Roth)^62^ into the ventral orbitofrontal cortex (OFC, AP: +4.2, ML: ±1.3, DV:-4.8; 1.0 µL/ hemisphere). GRABDA, iGluSnFr, and Cre (pENN.AAV_rg_-hSyn-HI-eGFP-Cre.WPRE.SV40, 105540-AAVrg; 0.5 µL at titer ≥ 7×10¹² vg/mL, deposited by James M Wilson) were combined and infused into the DMS of one hemisphere and the DLS of the other hemisphere (same coordinates as above). After viral infusion, optical fibers (Doric) were implanted over the DMS and DLS at the same location and secured with skull screws and cement, as above. This viral strategy allowed us to individually or jointly inhibit projections from the OFC (CNO, 3 mg/kg i.p., Hello Bio) and/or the IN Thal (salvinorin B, 15 mg/kg s.c., Hello Bio) to the DMS or DLS and perform photometry recordings from these same regions.

### Discrimination paradigm and strategy switching

Following the virus infusion surgery, rats first learned to generally nose-poke for a reward. They were trained to nose poke in either of two lit, active ports in an operant chamber to receive a sugar pellet reward (nose-poke pre-training, Fig S1a). As such, they all learned to use a simple visual cue to identify an active port. Once rats earned 50 sugar pellets in a 30-minute period, they went on to a second pre-training phase. For this phase, rats had to initiate each trial with a nose-poke on a central port, prior to nose-poking either of the two lit, active ports (Fig S1b). Once rats earned 50 sugar pellets in a 30-minute period in this second pre-training phase, the pre-training phases were complete. Rats that went on to be fast and slow switchers took a similar number of days to complete the two pre-training phases (Fig S1c). A side bias rating was taken on the final pre-training day, which was the number of nose-pokes on the preferred side divided by the total number of nose-pokes. Most rats had a side bias, but there were no differences between rats that went on to be fast or slow switchers (Fig S1d). Ten rats showed a right-side bias and 12 rats showed a left side bias. Rats then performed a complex discrimination paradigm where they continued to discriminate between the 2 active ports using a visual strategy (Fig. 1b). However, now they had to ignore the position of the visual cue, as the light cue appeared in either the left or right port, pseudorandomly. A trial began with a central port being illuminated. The rat had to nose-poke into this port to initiate the trial, similar to the second pre-training phase. Once the trial was initiated, rats were to respond into the port with the light regardless of where it was located. If they chose correctly, they again received a sugar pellet reward, if they chose incorrectly, they did not receive a pellet but there were no other programmed consequences. There was a brief inter-trial interval (4 seconds) and then the central port illuminated, indicating that a new trial was available. Rats did this for 100 trials/ day for 10 days. On the 11^th^ day the rats performed a strategy switching test. The test began with 20 “reminder” trials (pre-switch) that rewarded the same visual strategy that had been rewarded for the previous 10 days (Fig. 1c). On trial 21, the rewarded strategy switched to a response strategy (Fig. 1d). Thus, the rats had to learn to ignore the visual cue and respond either to the left or to the right port (response strategy). The rewarded port side was opposite each rat’s natural side bias as determined at the end of the pretraining phase. We quantified the number of trials and errors it took the rats to reach the performance criterion of 10 consecutive trials correct following the strategy switch (Fig. 1e).

### Photometry

DMS and DLS DA and Glut recordings were performed as we have reported in our prior studies^56^. For the first experiment, recordings occurred on discrimination days 1, 5, and 10, and during the strategy switching test. Data from the strategy switching test were divided into the first ∼30 trials after the strategy switch (early strategy switch), trials ∼60-90 (middle strategy switch), and the final ∼20 trials as rats reached the performance criterion (late strategy switch). For the second experiment, recordings occurred on discrimination training day 1,4,5, and 6. For the strategy switching test, we recorded the first 90 trials, split into 3 30-trial sets. During recording sessions rats were connected to a patch cable (200 μm diameter, Doric) attached to the fiber photometer (Plexon). The igluSnFR sensor (green) was excited at 465 nm with the minimum power that ensures a robust signal to noise ratio (80-150 μW) and that reduces photobleaching to prolong the duration of recording sessions (up to 45 min). The emission fluorescence of the sensor (525 nm) was acquired at a frequency of 30 Hz to allow sub-second time resolution using Plexon software. The GRABDA sensor (red) was excited with light at 560 nm and the emission (600 nm) was acquired with the same frequency. Emissions were collected simultaneously with a high quantum-efficiency CMOS camera. Glut and DA fluctuations were measured as the deflection in fluorescence relative to baseline levels (GuPPy^63^). Frames were interleaved with excitation at the isosbestic point (410 nm), which was then subtracted to control for movement artifacts. Relevant behaviors were time stamped to align to photometry signals. For the discrimination days, we time-stamped photometry measures to trial initiation, correct and incorrect choices. Pellet retrieval and no pellet were quantified relative to a correct and incorrect choice, respectively. For the strategy switching test we time-stamped the photometry signal to trial initiation, correct choices following the previous strategy, correct choices following the new strategy, and incorrect choices. Pellet retrieval and no pellet were quantified relative to a correct and incorrect choice. The photometry signal for each behavior was analyzed in different windows. For trial initiation, correct, and incorrect choices, we analyzed one second prior to the timestamp, for no pellet we analyzed the 1 second after choice, and for pellet retrieval we analyzed the 2 seconds while the rat is consuming the pellet.

### Cocaine self-administration (SA)

At least two days after the strategy switching test concluded, all rats were implanted with an indwelling intravenous catheter as we have previously published^18,56,58^. Following a 1-week recovery from catheter implant surgery, rats were trained to self-administer cocaine (0.5 mg/kg) paired with a tone/light cue such that each lever press on the active lever delivered both the cue and cocaine (FR1 schedule, Fig. 1f). The lever presses on the inactive lever had no programmed consequences. The sessions lasted 125 minutes. Once the rats took at least 15 infusions of cocaine with twice as many active as inactive lever presses for 2 consecutive days, the training criterion was reached, and rats began 10 days of cocaine self-administration (Fig. 1h). During these 10 days, rats either had access to cocaine continuously on an FR1 schedule for the entire 125-minute session (continuous access, first and second experiment), or they had access to cocaine in 5-minute bins every 30 minutes (intermittent access, first experiment only). Intermittent access schedules have been shown to precipitate a more punishment resistant phenotype^18,21,22^; thus, this study design allowed us to examine how strategy switching performance interacts with cocaine access schedule. On the 11^th^ day, all rats were given continuous access to cocaine for the entire 125-minute session (continuous access test) to more directly compare cocaine intake between the two access schedules. On day 12, all rats performed a punishment resistance test (Fig. 1k), where cocaine infusions were first earned on an FR1 schedule for 20 minutes with no shock, followed by 90 minutes with shock. During the shock phase, cocaine infusions were paired with a foot shock that increased in intensity every 10 minutes on a tenth-log_10_ scale from 0.13 to 0.79 milliamps^18–20^. Cocaine infusions across the test were binned into four groups, no shock (20 minutes), low shock (0.13-0.2 mA, 30 minutes), medium shock (0.25-0.4 mA, 30 minutes), and high shock (0.5-0.79 mA, 30 minutes).

### Histology

Rats were euthanized, perfused transcardially with 4% paraformaldahyde, and decapitated. Brains slices were harvested and mounted for immunofluorescent confirmation of viral placement and spread and photometry fiber placement. An anti-mCherry-Rabbit monoclonal recombinant IgG (Sysy, Catalog no. 409008, 1:500) and a goat anti-rabbit secondary (abcam150077, 1:500) were used for confirmation of the GRABDA. An anti-GFP-chicken monoclonal recombinant IgG (abcam 13970, 1:2000) and a donkey anti-chicken secondary (Jackson Immuno Research Labs; Catalog no. NC0456003 via Fisher, 1:5000) were used for confirmation of the igluSnFR.

### Data Analysis

To examine how photometry signaling in the DMS and DLS relates to strategy switching performance and subsequent cocaine intake and punishment resistance, we analyzed our data both as a continuum and using a median split to parse groups into “fast” and “slow” switchers. Behavioral data (e.g., trials, errors, cocaine infusions, etc.) was analyzed by multi-factor ANOVA with repeated measures as appropriate. When analyzing the data categorically, we used strategy switching errors to perform a median split into “fast” and “slow” strategy switchers. Significant interactions were followed by Tukey’s post-hoc test with p<0.05 considered significant. Fiber photometry data were processed in real-time to collect z-scores and a heat map. Behavioral data were collected by MedPC and TTL signals were sent to Plexon to ensure data were within the identical “time-space.” We compared areas under the curve (AUC) across a 1-2 second time window for each behavioral time stamp using the same analysis strategy as above. Because we measured DMS and DLS DA and Glut in every rat, we also analyzed how these signals affected each other within and between rats using repeated measures ANOVA, multiple linear regression models, and correlation matrices. Note that p values in correlation matrices are not corrected for multiple comparisons. We performed our first experiment with 23 rats. One rat was removed from all analyses due to a dislodged headcap. Five rats were removed from the cocaine analysis due to catheter patency issues, and one was removed for errors made administering the punishment sensitivity/ resistance test. The second experiment was performed with 15 rats. One rat was removed from all analyses due to injection errors. Two of seven rats were not used for figure 8a because 1 rat finished the strategy switching test prior to the CNO injection and 1 rat had a medial placement in the OFC. Two rats were removed from the cocaine analysis because they did not meet the self-administration acquisition criteria, and one was removed because the shock floor malfunctioned during the punishment sensitivity/ resistance test.

Linear Discriminant Analysis (LDA) was used to predict two behavioral metrics: switching speed and punishment sensitivity, based on neurotransmitter activity metrics, which served as predictor variables. To identify the most effective combinations of predictors, separate binary classifiers were trained for each behavioral metric using all possible combinations of 1 to 14 predictor variables. 8-fold cross-validation was used to measure the classification accuracy of each model. Models with an average classification accuracy of 100% were then examined to identify the specific predictor variables that were used. The predictive strength of each individual variable was further assessed by ranking them according to the negative logarithm of the p-value obtained from a Chi-square test. This test evaluated each variable’s ability to discriminate between the two categories of each behavioral metric.

### Strategy switching categorization for experiment 2

We aimed to measure the effects of inhibiting OFC and IN Thal inputs to the dorsal striatum on Glut and DA release and behavior. However, based on the results from our first experiment, we hypothesized that inhibiting these projections may result in different effects in rats that would be fast or slow strategy switchers. However, because we manipulated projections during the strategy switching test, we could not categorize them based on their actual performance. Our linear discriminant analysis model identified hundreds of combinations of the neurotransmitter metrics that yielded perfect classification of switching speed. Thus, we selected 3 3-metric models that used metrics from early in the discrimination paradigm (day 1 and 5) and early in the strategy switching test, i.e. when the rats were unmanipulated (vehicle condition). Each model categorized rats as fast if they featured 2 out of 3 of the following: Glut AUC measures >2.5, DA AUC measures <0, and Glut/DA diff AUC measures > 2.5. They were categorized as slow if they were the opposite of that for 2 out of 3 measures. For rats where the different models disagreed, the final categorization was based on the majority opinion, 2 out of 3 models (see Fig S5).

---

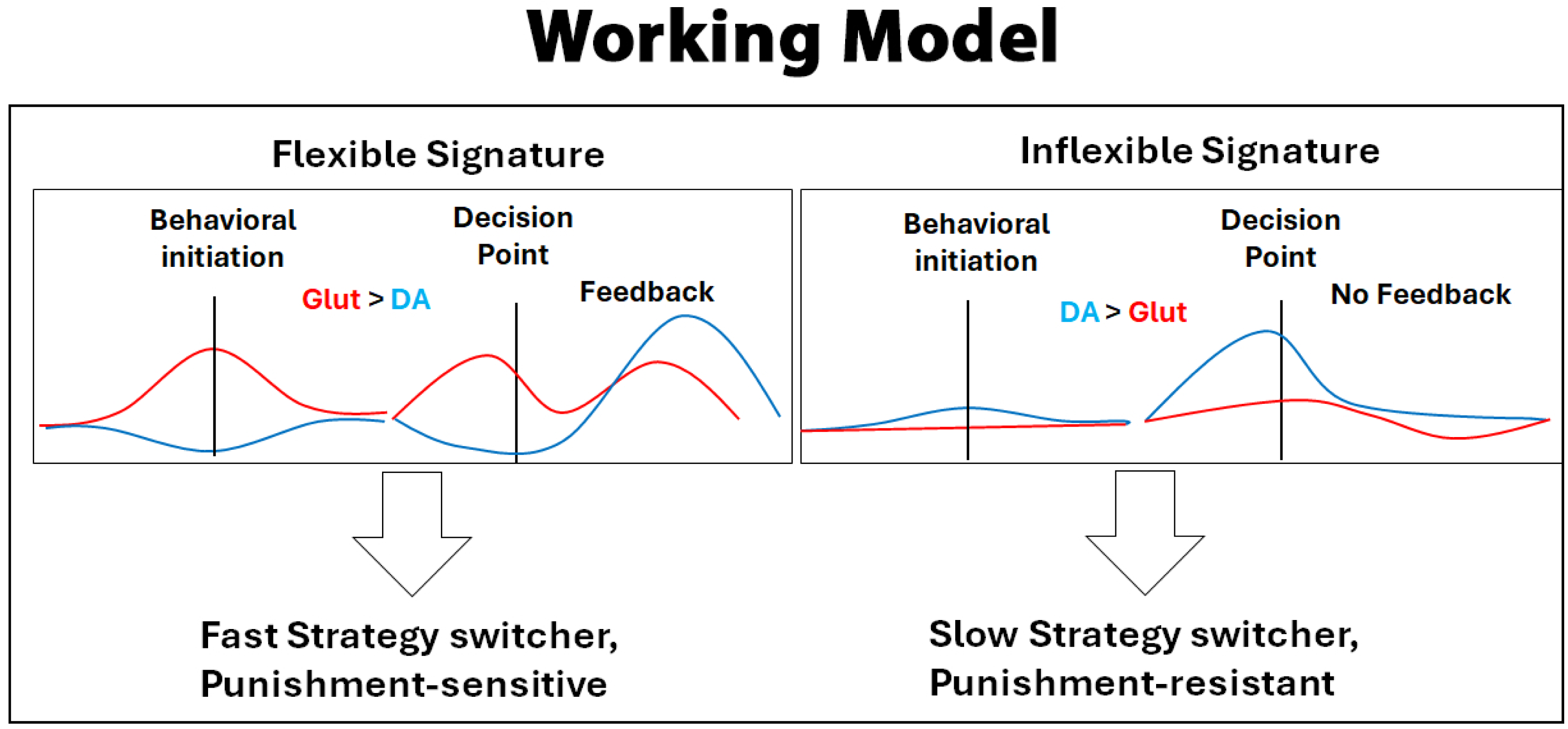

## Notes

### Competing Interest Statement

The authors have declared no competing interest.

### Summary of Updates

Manuscript edited down and figures edited.

