## Supplemental figure legends for "Dynamic, behavior-dependent interactions between dorsal striatal dopamine and glutamate release predict cognitive flexibility and punishment resistant cocaine use"

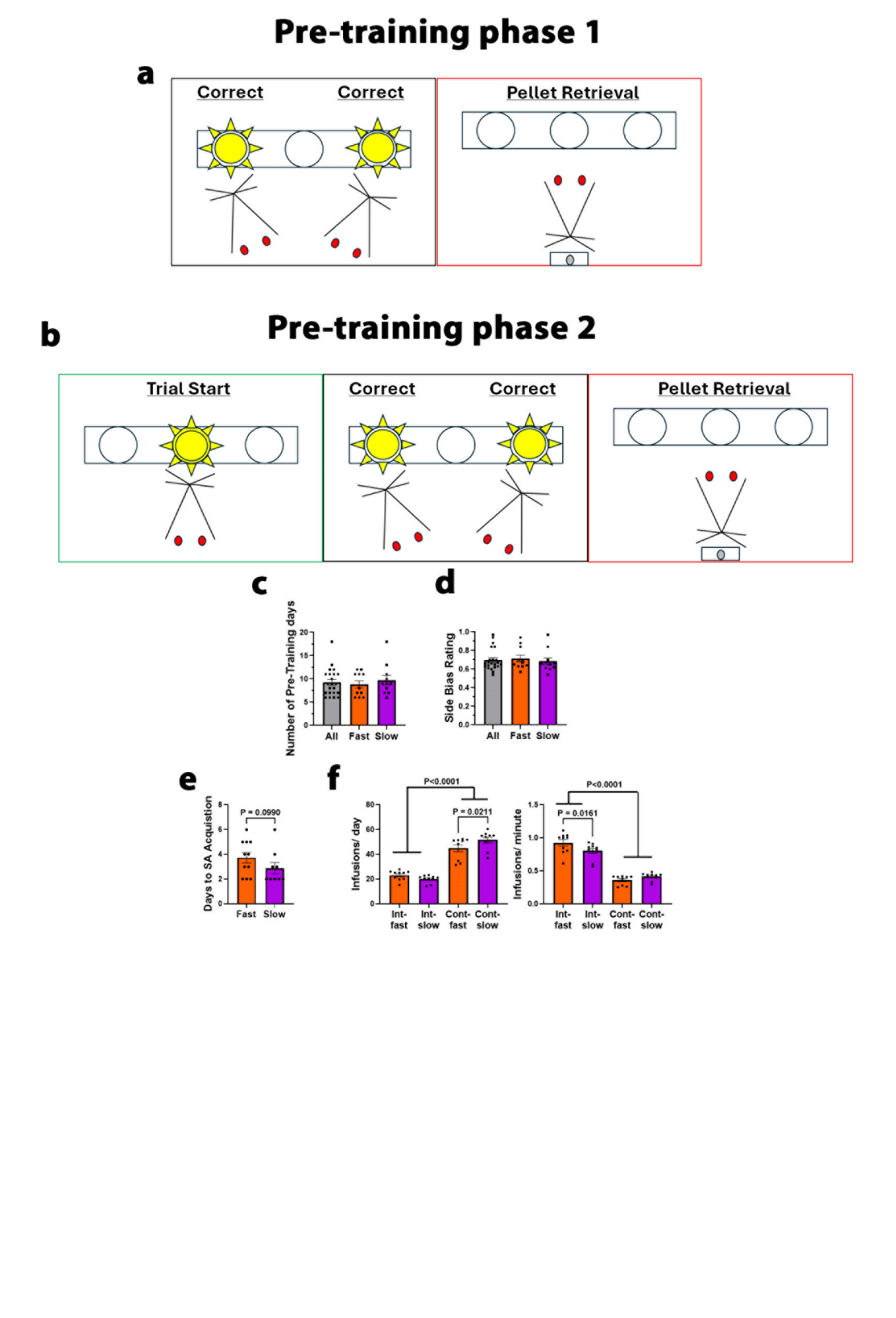

**Figure S1 Fast and slow switchers were not different in number of pre-training days or presence of a side bias: a)** Pre-training phase 1 entails the rat learning to nose-poke either of 2 active ports to receive a sugar pellet. The active ports are indicated by a lit port light. They continue this phase until they earn 50 pellets in 30 minutes, and this typically takes 5-7 days. **b)** Pre-training phase 2 is the same as 1 except the rat must now initiate the trial by nose-poking a central port. Once the rat nose-pokes the central port, the central port light goes off and the 2 active port lights turn on. They continue this phase until they again earn 50 pellets in 30 minutes, and this typically takes 1-3 days. **c)** Rats that went on to be fast or slow switchers were not different in the number of pre-training days they took to move on to the discrimination phase. **d)** Rats that went on to be fast or slow switchers were not different in the degree to which they naturally preferred one port or the other (side bias). **e)** Slow switchers (2.9±0.5 days) reached self-administration criterion in fewer days than fast switchers (3.7±0.4 days; t(18)=1.34, P=0.099). **f)** Continuous access rats took more cocaine infusions per day (fast switchers: 44.9±2.7, slow switchers: 51.6±*2.*3) relative to intermittent access rats (fast switchers: 23.1±1.2, slow switchers: 20.1±1.0; P<0.0001) due to their continuous access, but intermittent access rats took more cocaine per minute (fast switchers: 0.9±0.05, slow switchers: 0.8±0.04) relative to continuous access rats (fast switchers: 0.4±0.02, slow switchers: 0.4±0.02; P<0.0001). Similar to previous studies, fast switchers with continuous access took fewer cocaine infusions per day than slow switchers (P=0.02), but fast switchers with intermittent access took more cocaine infusions/ minute (P=0.02).

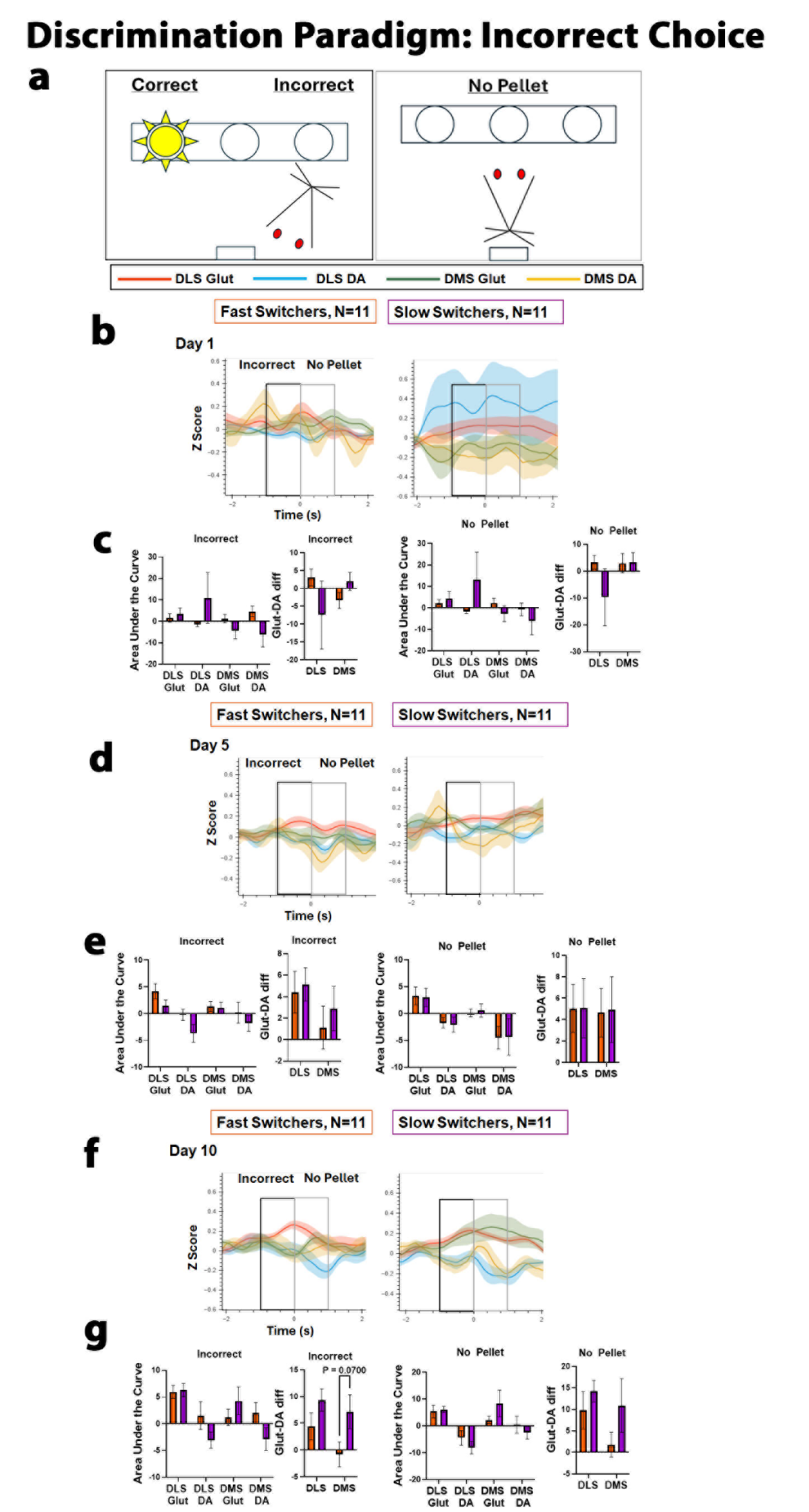

**Figure S2 Fast and Slow switchers were not different prior to an incorrect choice or when no pellet is received during Discrimination paradigm: a)** This figure depicts mean tracings aligned to incorrect choices and subsequent no pellet retrieval of DLS Glut (red), DLS DA (blue), DMS Glut (green), and DMS DA (yellow) release across fast (left, N=11) and slow (right, N=11) switchers. Area under the curve is shown as in prior figures. **b)** Tracings were highly variable with no significant differences between fast and slow switchers. **c)** AUC Mean ± SEM for day 1 are as follows: Incorrect: DLS Glut (fast: 1.7 ± 2.0, slow: 3.4 ± 2.7) DLS DA (fast: -1.4 ± 1.0, slow: 10.9 ± 11.9) | DMS Glut (fast: 1.2 ± 2.0, slow: -4.2 ± 3.8) DMS DA (fast: 4.6 ± 2.5, slow: -6.2 ± 5.8). No Pellet: DLS Glut (fast: 2.1 ± 1.8, slow: 4.2 ± 3.4) DLS DA (fast: -1.7 ± 0.9, slow: 13.1 ± 13.0) | DMS Glut (fast: 2.3 ± 2.2, slow: -2.7 ± 3.7) DMS DA (fast: -0.7 ± 3.0, slow: -6.1 ± 6.4). **d)** There were no differences on day 5. **e)** Incorrect: DLS Glut (fast: 4.2 ± 1.4, slow: 1.4 ± 1.2) DLS DA (fast: -0.2 ± 1.0, slow: -3.7 ± 1.6) | DMS Glut (fast: 1.3 ± 0.9, slow: 1.1 ± 1.0) DMS DA (fast: 0.2 ± 1.9, slow: -1.8 ± 1.5). No Pellet: DLS Glut (fast: 3.3 ± 1.7, slow: 3.0 ± 1.7) DLS DA (fast: -1.7 ± 1.0, slow: -2.0 ± 1.4) | DMS Glut (fast: 0.1 ± 0.7, slow: 0.6 ± 1.2) DMS DA (fast: -4.5 ± 2.1, slow: -4.3 ± 3.4). **f)** There were no differences on day 10. **g)** Incorrect: DLS Glut (fast: 6.0 ± 1.2, slow: 6.3 ± 1.2) DLS DA (fast: 1.5 ± 2.6, slow: 6.3 ± 1.2 | DMS Glut (fast: 1.2 ± 1.5, slow: 4.3 ± 2.6) DMS DA (fast: 2.0 ± 1.9, slow: -2.9 ± 2.1). No Pellet: DLS Glut (fast: 5.3 ± 2.4, slow: 6.2 ± 1.1) DLS DA (fast: -4.4 ± 2.8, slow: -8.12 ± 2.4) | DMS Glut (fast: 2.2 ± 1.4, slow: 8.4 ± 5.0) DMS DA (fast: 0.4 ± 3.2, slow: -2.5 ± 2.4).

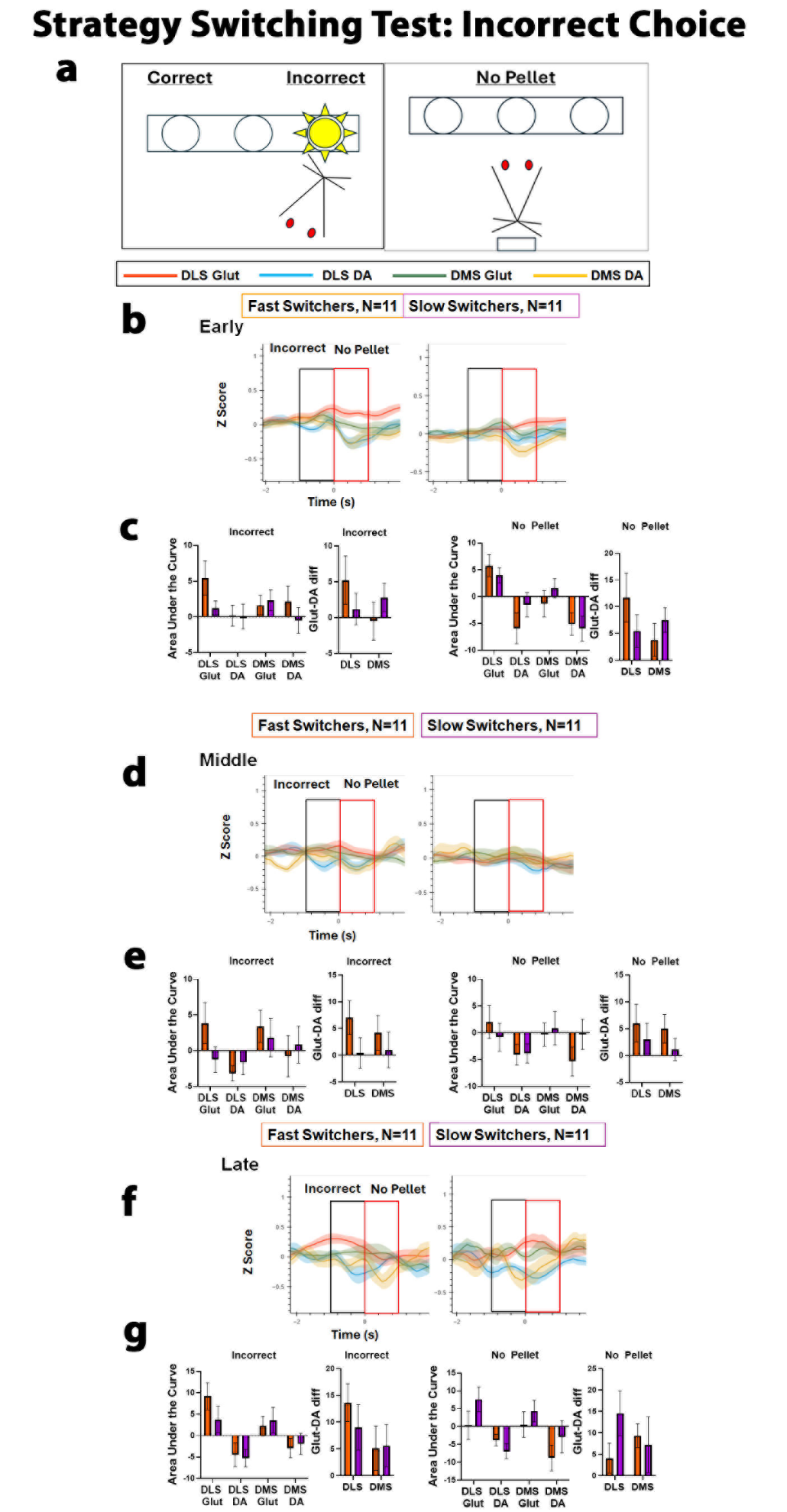

**Figure S3** **Fast and Slow switchers were not different prior to an incorrect choice or when no pellet is received during the strategy switching test: a)** This figure depicts mean tracings aligned to incorrect choices and subsequent no pellet retrieval of DLS Glut (red), DLS DA (blue), DMS Glut (green), and DMS DA (yellow) release across fast (left, N=11) and slow (right, N=11) switchers. Area under the curve is shown as in prior figures. **b)** Tracings indicated no differences between fast and slow switchers. **c)** AUC Mean ± SEM for early in the strategy switching test are as follows: Incorrect: DLS Glut (fast: 5.4 ± 2.4, slow: 1.2 ± 1.0) DLS DA (fast: 0.2 ± 1.5, slow: 0.03 ± 1.7) | DMS Glut (fast: 1.6 ± 1.4, slow: 2.3 ± 1.5) DMS DA (fast: 2.1 ± 2.2, slow: -0.5 ± 1.8). No Pellet: DLS Glut (fast: 5.8 ± 2.1, slow: 4.0 ± 1.4) DLS DA (fast: -6.0 ± 2.9, slow: -1.5 ± 2.3) | DMS Glut (fast: -1.3 ± 2.5, slow: 1.6 ± 1.8) DMS DA (fast: -5.1 ± 2.1 0, slow: -5.9 ± 2.3). **d)** There were no differences during the middle portion of the test. **e)** Incorrect: DLS Glut (fast: 3.8 ± 2.9, slow: -1.3 ± 1.8) DLS DA (fast: -3.2 ± 1.1, slow: -1.7 ± 1.7) | DMS Glut (fast: 3.4 ± 2.3, slow: 1.8 ± 2.7) DMS DA (fast: -0.8 ± 2.9, slow: 0.9 ± 2.6). No Pellet: DLS Glut (fast: 2.0 ± 3.1, slow: -0.8 ± 2.6) DLS DA (fast: -4.0 ± 2.0, slow: -3.8 ± 1.8) | DMS Glut (fast: -0.3 ± 2.2, slow: 0.9 ± 3.1) DMS DA (fast: -5.3 ± 2.7, slow: -0.3 ± 2.8). **f)** There were no differences on day 10. **g)** Incorrect: DLS Glut (fast: 9.2 ± 3.1, slow: 3.8 ± 3.0) DLS DA (fast: -4.4 ± 2.8, slow: -5.2 ± 2.1) | DMS Glut (fast: 2.2 ± 2.3, slow: 3.6 ± 3.0) DMS DA (fast: -2.9 ± 2.2, slow: -2.0 ± 2.5). No Pellet: DLS Glut (fast: 0.3 ± 4.0, slow: 7.7 ± 3.5) DLS DA (fast: -3.8 ± 1.6, slow: -6.9 ± 2.1) | DMS Glut (fast: 0.6 ± 3.6, slow: 4.4 ± 3.0) DMS DA (fast: -8.8 ± 3.6, slow: -2.8 ± 4.5).

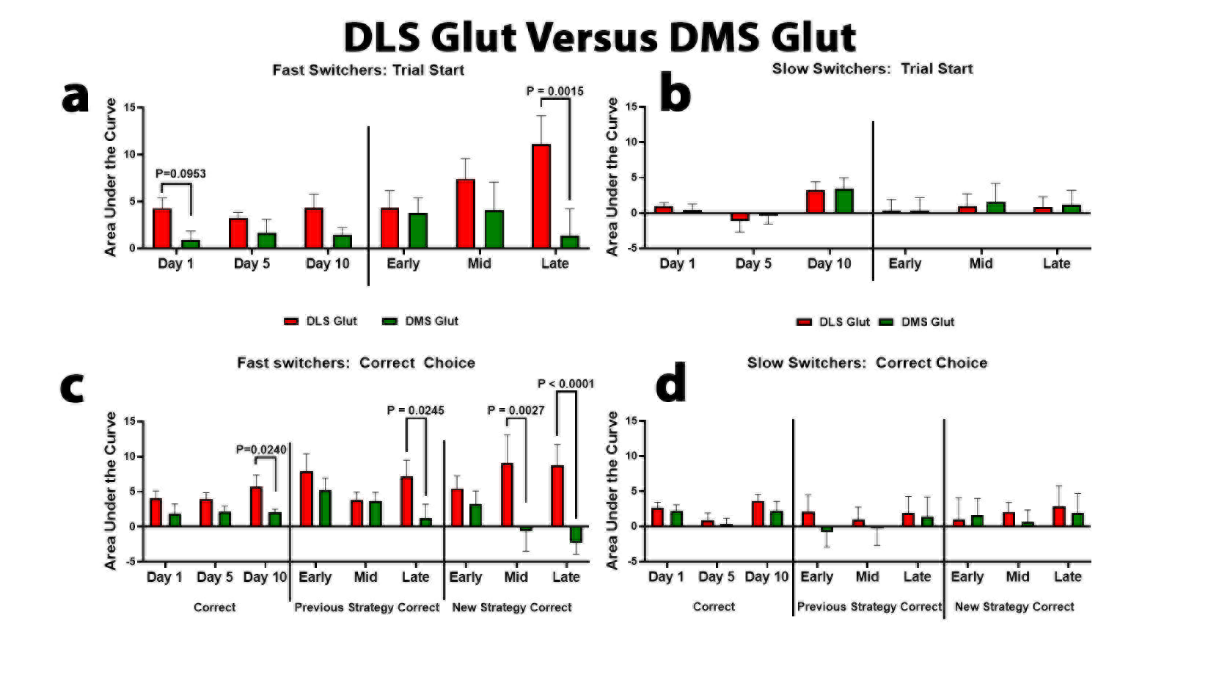

**Figure S4 DLS and DMS Glut release changes both converge and diverge across the discrimination paradigm and strategy switching test:** We analyzed DLS and DMS Glut release across time at the two behavioral timestamps that produced the effects described above: trial start and correct choice. **a)** At trial start there was a main effect of dorsal striatum region (F(1,20)=6.40, P=0.0199), such that DLS Glut release was greater than DMS Glut release throughout performance of the discrimination paradigm and into the early portion of the strategy switching test. DLS Glut increased throughout the strategy switching test reaching a peak late in the test. Alternatively, DMS Glut peaked during the early and middle portions of the strategy switching test, resulting in a difference between DLS and DMS Glut on day 1 of the discrimination paradigm (P=0.0953) and late in the strategy switching test (P=0.0015), but not at other time points. **b)** DMS and DLS Glut release remained low and even in slow switchers across all time points except at day 10. Interestingly, the release ratio of DLS and DMS Glut on day 10 in the slow switchers resembles the ratio of fast switchers early in the strategy switching test. **c)** Prior to a correct choice, there was a main effect of dorsal striatum region (F(1,20)=15.3, P=0.0009) and a marginal interaction between region and time (F(8,150)=1.96, p=0.055; Fig. 6c). DLS Glut, but not DMS Glut, increased prior to a correct choice across performance of the discrimination paradigm, such that DLS Glut was greater than DMS Glut by day 10 (P=0.024). This difference was no longer present early in the strategy switching test when making a correct choice using the previous strategy as Glut release increased in both regions, but particularly in the DMS. Uniform release in both the DMS and DLS was maintained during the middle portion of the test at this timestamp but then separated again late in the test (P=0.0245) as DLS Glut increased and DMS Glut decreased. When making a correct choice following the new strategy, there was a different pattern. During the early portion of the test, the difference between DLS and DMS Glut seen on day 10 went away, and this was again due to an increase in DMS Glut. However, there was a quick re-emergence of the separation of DMS and DLS Glut release as DLS Glut increased and DMS Glut decreased during the middle of the test (P=0.0027) and then became further pronounced late in the test (p<0.0001). **d)** These patterns were absent in the slow switchers, who despite having similar release measures across the discrimination paradigm to fast switchers, featured a surprising reduction in both DMS and DLS Glut early in the strategy switching test relative to their day 10 release levels. Thus, it appears that in fast switchers, DMS Glut release is observed when contingencies change and the action-outcome relationship becomes uncertain, while DLS Glut dominates when actions are well-learned and outcomes are certain, whereas slow switchers fail to engage these striatal signaling dynamics under changing conditions.

The final experiment may also help in disentangling these views to some extent. We showed that OFC inhibition reduced DMS and DLS Glut prior to a correct choice using the previous strategy but only reduced DLS Glut at trial start. Alternatively, IN Thal inhibition reduced both DLS and DMS Glut at trial start, but only DLS Glut prior to a correct choice using the previous strategy. Therefore, one possibility is that DLS and DMS Glut inputs from OFC may be working cooperatively at choice points with high uncertainty, but competitively at behavioral initiation with low uncertainty. Likewise, DLS and DMS Glut inputs from IN Thal may be working cooperatively at behavioral initiation when uncertainty is high and competitively at choice points when uncertainty is low.

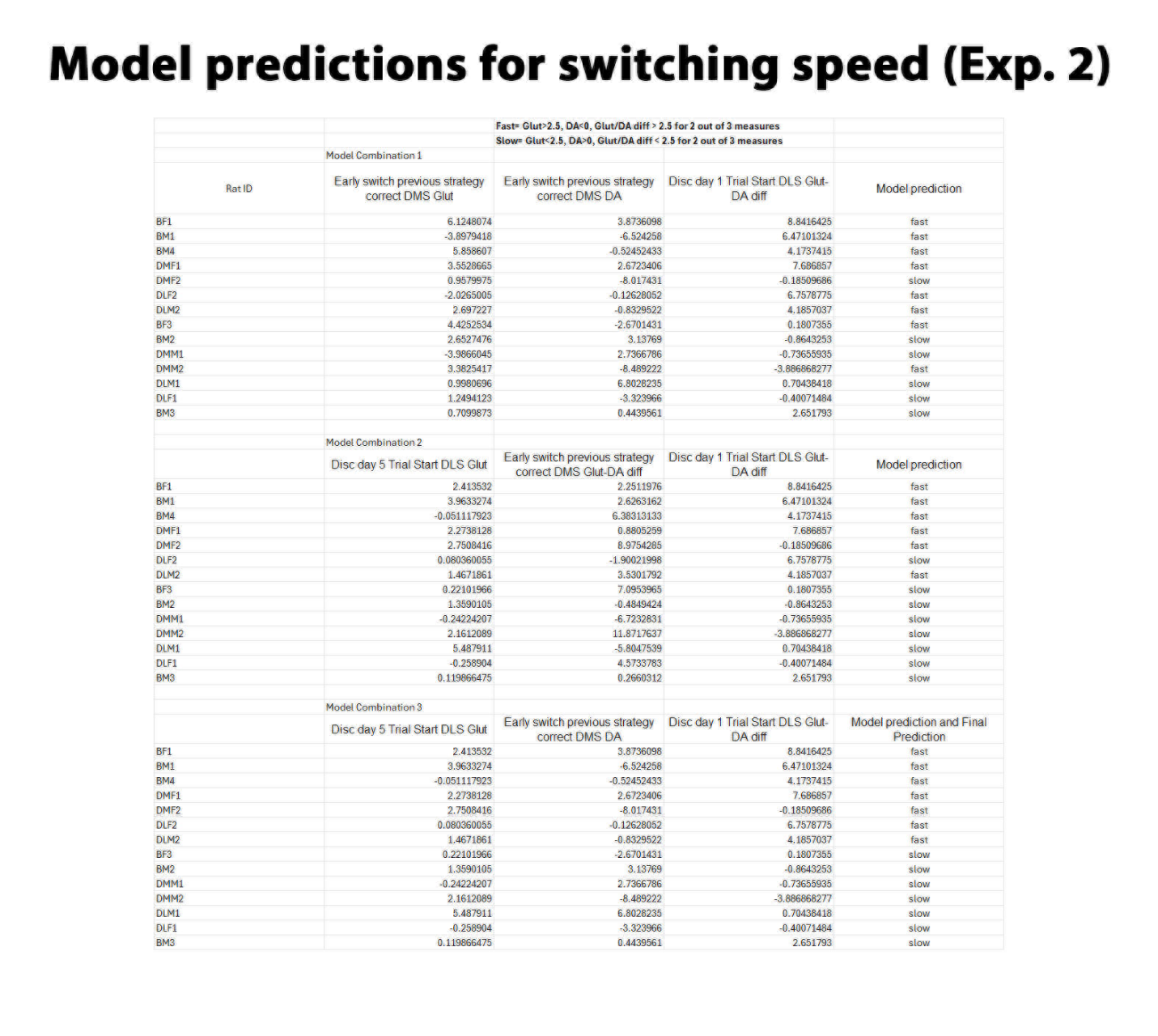

**Fig S5 Model predictions for switching speed for experiment 2:** Our linear discriminant analysis model identified hundreds of combinations of the neurotransmitter metrics that yielded perfect classification of switching speed. Thus, we selected 3 3-metric models that used metrics from early in the discrimination paradigm (day 1 and 5) and early in the strategy switching test, i.e. when the rats were unmanipulated (vehicle condition). Each model categorized rats as fast if they featured 2 out of 3 of the following: Glut AUC measures >2.5, DA AUC measures <0, and Glut/DA diff AUC measures > 2.5. They were categorized as slow if they were the opposite of that for 2 out of 3 measures. Rat ID is on the left column; the next 3 columns include the neurotransmitter metric and the raw AUC values for each rat. The final column indicates the model’s “decision” on whether the rat would have been “fast” or “slow”. For rats where the different models disagreed, the final categorization was based on the majority opinion, 2 out of 3 models. Model 3 shows the decision for that model, which coincides with the majority opinion and final categorization for each rat. Seven rats were categorized as fast switchers, 5 were categorized as fast by all 3 models and 2 were categorized as fast by 2 out of 3 models. One additional rat was categorized as fast by the model (BF2) but was removed from all analysis due to injection issues during the strategy switching test. Seven rats were categorized as slow switchers. Again, 5 rats were categorized as slow switchers by all 3 models and 2 were categorized as slow switchers by 2 out of the 3 models.

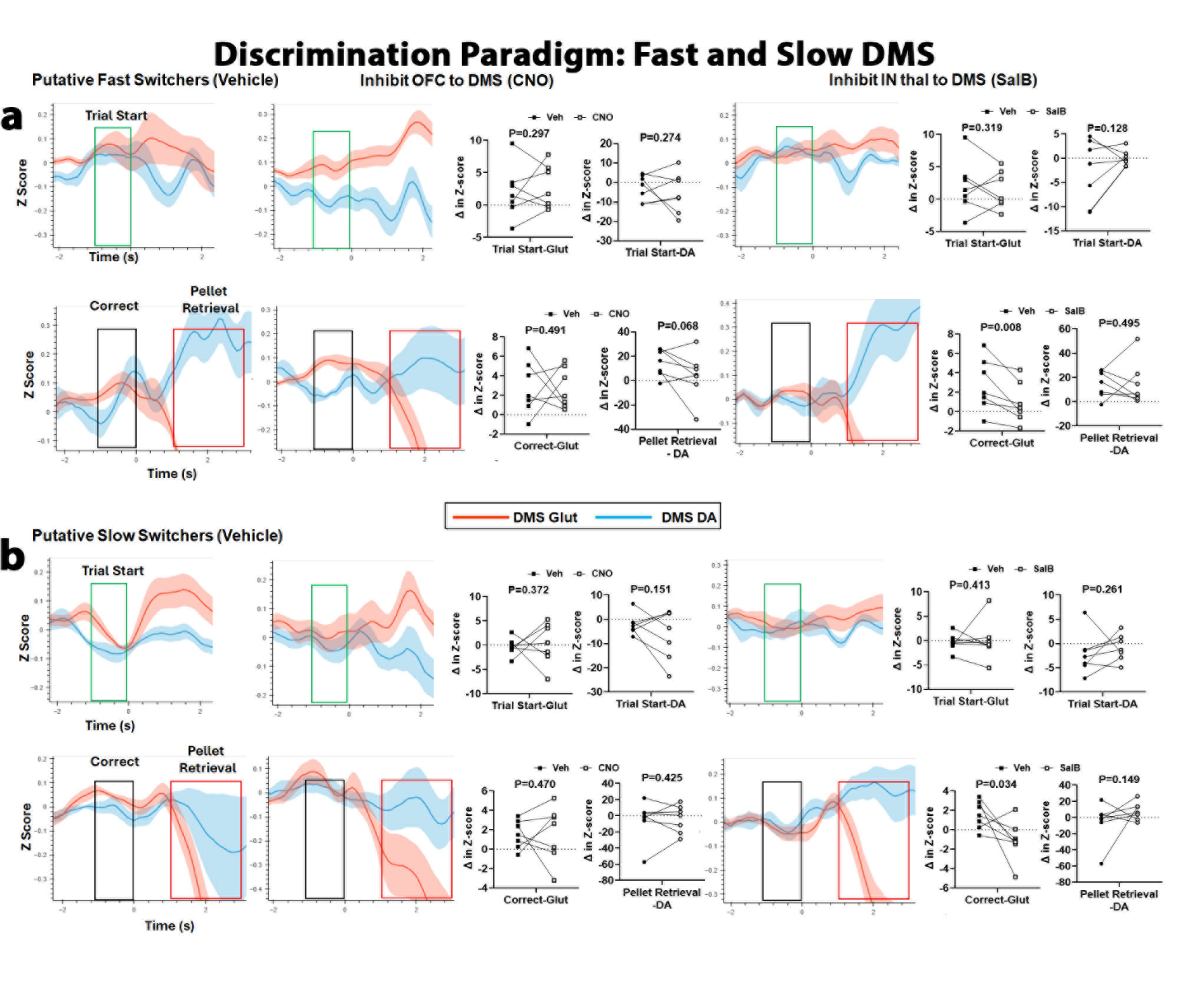

**Figure S6** **Inhibition of IN Thal inputs during Discrimination paradigm reduce DMS Glut prior to a correct choice in both putative fast and slow switchers:** Mean tracings are aligned to trial start (green box), correct choices (black box), and subsequent pellet retrieval (red box) of DMS Glut (red) and DLS DA (blue). Data across rows are within subjects and represents the change in signal (Z-score) from no manipulation (vehicle, left) to inhibition of OFC (CNO, middle) and then inhibition of IN Thal (SalB, right) inputs to dorsal striatum. The tracings are quantified using area under the curve (Mean ± SEM) for the 1 or 2 seconds in the colored box, as indicated. **a) Fast switchers:** Inhibition of OFC inputs did not affect DMS Glut or DA at Trial start (Glut- Veh: 1.99 ± 1.54, CNO: 2.80 ± 1.28, SalB: 1.37 ± 1.10; DA- Veh: -2.71 ± 2.50, CNO: -5.40 ± 3.96, SalB: 0.003 ± 0.63). OFC inhibition also did not affect DMS Glut prior to a correct choice (Glut-Veh: 2.75 ± 1.02, CNO: 2.72 ± 0.78) but did reduce DMS DA release during pellet retrieval (DA- Veh: 14.74 ± 4.17, CNO: 3.82 ± 7.27). Inhibition of IN Thal inputs reduced Glut release prior to a correct choice (Glut-0.90 ± 0.79) but did not affect DA (14.84 ± 6.81). **b) Slow switchers:** Inhibition of OFC or IN Thal had no effect on Glut or DA release at trial start (Glut-Veh: -0.32 ± 0.67, CNO: 0.39 ± 1.63, SalB: 0.04 ± 1.56; DA- Veh: -2.12 ± 1.61, CNO: -6.34 ± 3.90, SalB: -0.84 ± 1.05). Inhibition of OFC also had no effect prior to a correct choice, but IN Thal inhibition reduced Glut release, similar to the fast switchers (Correct-Glut-Veh: 1.52 ± 0.55, CNO: 1.61 ± 1.09, SalB: -1.08 ± 0.79; DA- Veh: -5.28 ± 9.29, CNO: -3.99 ± 6.46, SalB: 7.86 ± 4.19).

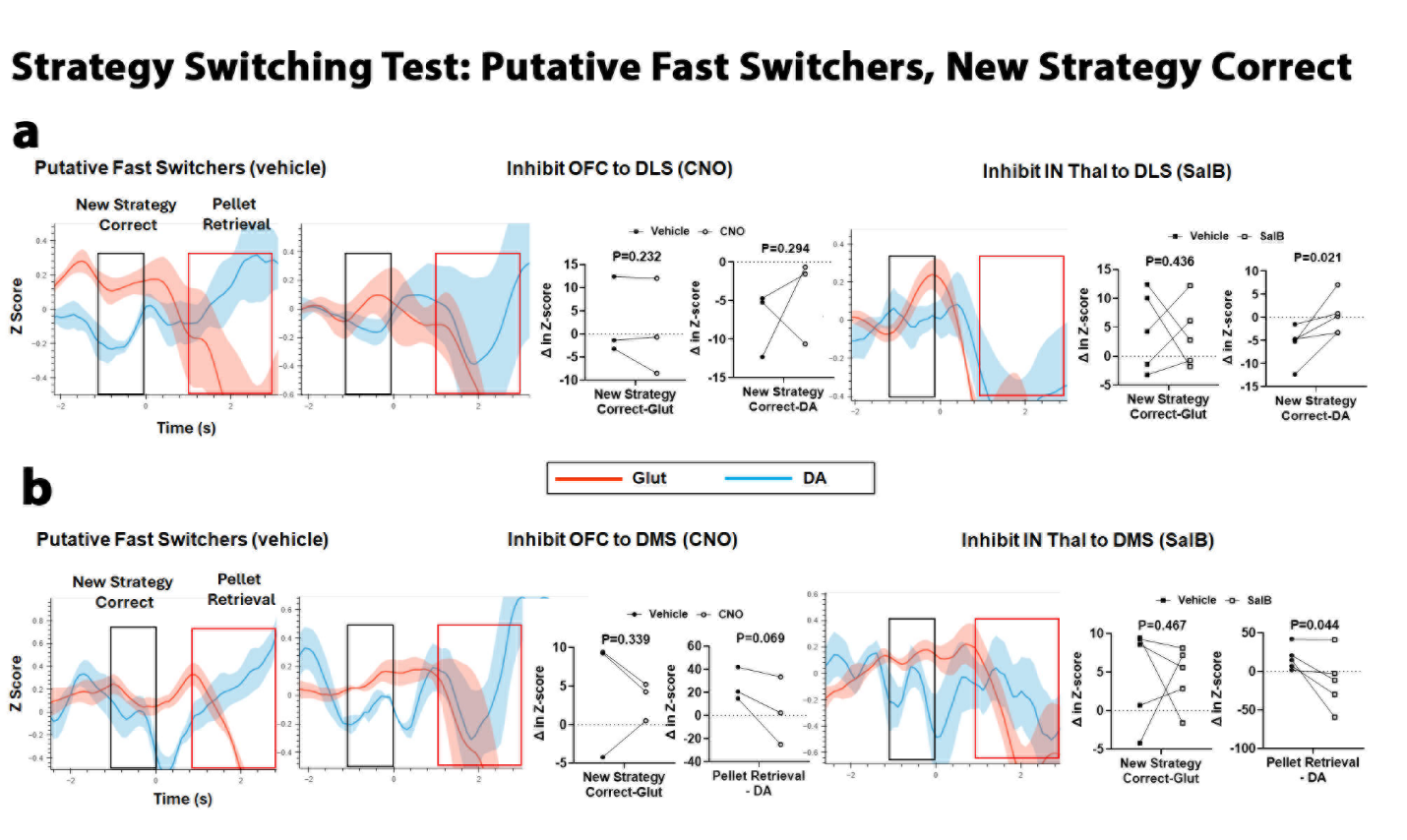

**Figure S7 Inhibition of OFC and IN Thal inputs reduce DMS DA during pellet retrieval and inhibition of IN Thal increases DLS DA prior to a correct choice in fast switchers prior to a new strategy correct choice:** Mean tracings are aligned to correct choices following the new strategy (black box), and subsequent pellet retrieval (red box) of Glut (red) and DA (blue). Data across rows are within subjects and represents the change in signal (Z-score) from no manipulation (vehicle, left) to inhibition of OFC (CNO, middle) and then inhibition of IN Thal (SalB, right) inputs to dorsal striatum. The tracings are quantified using area under the curve (Mean ± SEM) for the 1 or 2 seconds in the colored box, as indicated. **a) DLS:** Inhibition of OFC did not affect Glut or DA prior to a correct choice (Glut- Veh: 2.62 ± 4.94, CNO: 0.97 ± 5.98; DA- Veh: -7.43 ± 2.46, CNO: -4.28 ± 3.19). Inhibit IN Thal also did not affect Glut (Veh: 4.45 ± 3.07, SalB: 3.72 ± 2.54) but did result in an increase in DA (Veh: -5.72 ± 1.78, SalB: 0.32 ± 1.90). **b) DMS:** Inhibition of OFC to DMS did not affect Glut prior to a correct choice (Veh: 4.83 ± 4.53, CNO: 3.33 ± 1.44) but did result in a reduction in DA during pellet retrieval (Veh: 25.81 ± 8.23, CNO: 3.55 ± 16.90). Inhibit of IN Thal to DMS produced the same effect (Glut-Veh: 4.75 ± 2.78, SalB: 4.43 ± 1.76; DA- Veh: 17.27 ± 6.96, SalB: -12.49 ± 16.57).

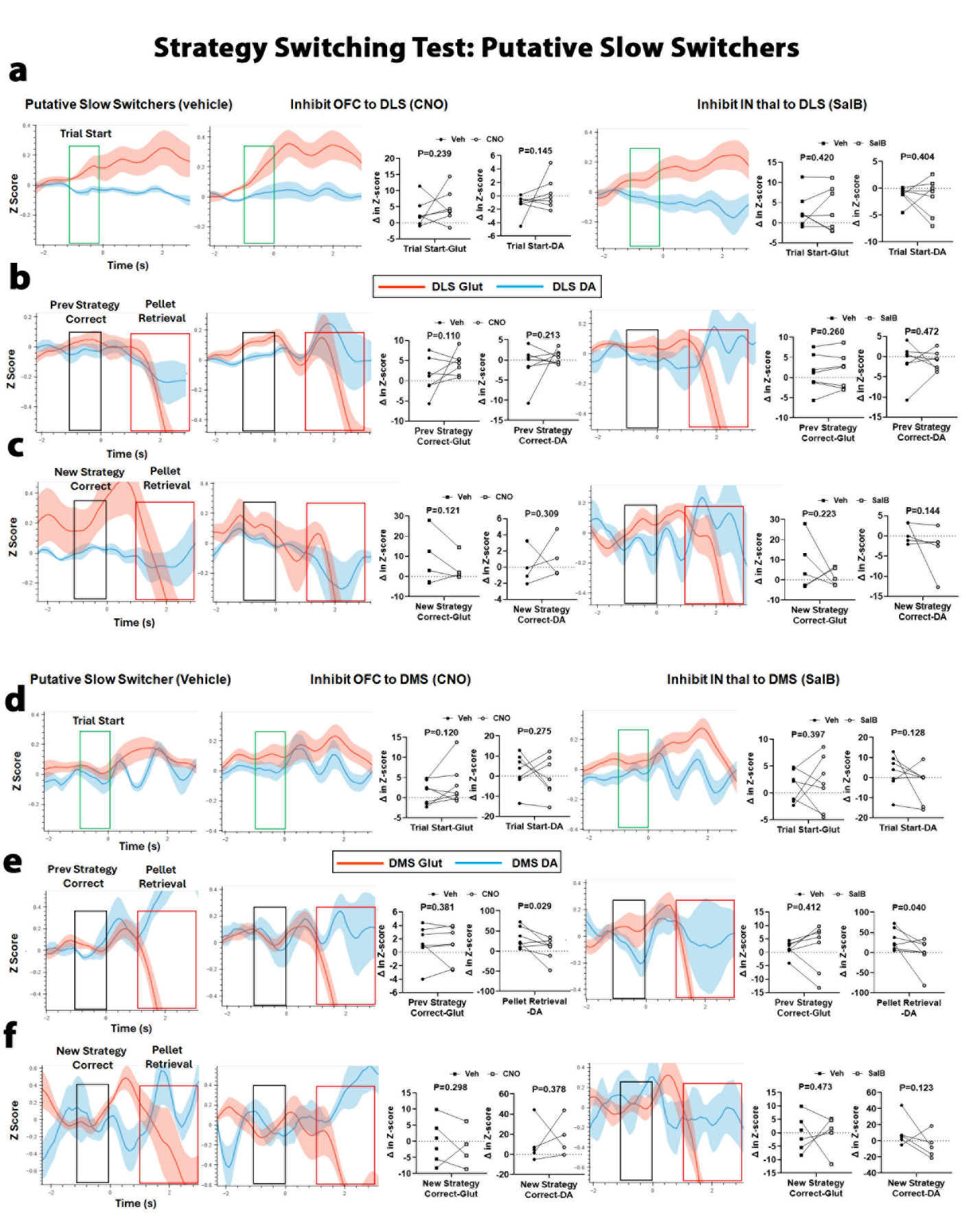

**Figure S8 Inhibition of OFC and IN Thal inputs have no effect on DLS or DMS Glut at any behavioral timestamp during the strategy switching test in putative slow switchers but do reduce DMS DA at pellet retrieval:** Mean tracings are aligned to trial start (green box), correct choices (black box), and subsequent pellet retrieval (red box) of region-specific Glut (red) and DA (blue) release. Data across rows are within subjects and represents the change in signal (Z-score) from no manipulation (vehicle, left) to inhibition of OFC (CNO, middle) and then inhibition of IN Thal (SalB, right) inputs to dorsal striatum. The tracings are quantified using area under the curve (Mean ± SEM) for the 1 or 2 seconds in the colored box, as indicated. **DLS: a)** Inhibition of OFC and IN Thal has no effect on Glut or DA in the DLS at trial start (Glut-Veh: 3.06 ± 1.58, CNO: 5.08 ± 1.94, SalB: 3.38 ± 2.07; DA- Veh: -1.23 ± 0.58, CNO: 0.37 ± 0.90, SalB: -1.60 ± 1.32). **b)** Inhibition of OFC and IN Thal has no effect on Glut or DA in the DLS prior to a correct choice following the previous strategy (Glut-Veh: 1.21 ± 1.69, CNO: 4.26 ± 1.07, SalB: 1.61 ± 1.67; DA- Veh: -1.16 ± 1.77, CNO: 0.75 ± 0.62, SalB: -1.03 ± 0.88). **c)** Inhibition of OFC and IN Thal has no effect on Glut or DA in the DLS prior to a correct choice following the new strategy (Glut-Veh: 7.52 ± 5.82, CNO: 4.28 ± 3.42, SalB: 1.56 ± 1.98; DA- Veh: 0.64 ± 1.11, CNO: 1.09 ± 1.29, SalB: -3.13 ± 2.54). **DMS: d)** Inhibition of OFC and IN Thal has no effect on Glut or DA in the DMS at trial start (Glut-Veh: 1.28 ± 1.11, CNO: 3.32 ± 1.93, SalB: 1.86 ± 1.88; DA- Veh: 2.53 ± 3.32, CNO: -0.32 ± 3.76, SalB: -3.00 ± 3.40). **e)** Inhibition of OFC and IN Thal has no effect on Glut in the DMS prior to a correct choice following the previous strategy (Veh: 1.35 ± 1.03, CNO: 1.14 ± 1.04, SalB: 2.00 ± 3.35) but does result in a reduction in DMS DA during pellet retrieval (Veh: 32.48 ± 9.80, CNO: 5.83 ± 10.42, SalB: -1.37 ± 14.45). **f)** Inhibition of OFC and IN Thal has no effect on Glut or DA in the DMS prior to a correct choice following the new strategy (Glut-Veh: -0.20 ± 2.70, CNO: -1.95 ± 3.14, SalB: -0.07 ± 3.08; DA- Veh: 9.79 ± 7.14, CNO: 17.54 ± 9.69, SalB: -5.85 ± 6.96).
